# Association between breastfeeding during infancy and white matter microstructure in early childhood

**DOI:** 10.1101/2021.01.05.425482

**Authors:** Preeti Kar, Jess E. Reynolds, Melody N. Grohs, Rhonda C. Bell, Megan Jarman, Deborah Dewey, Catherine Lebel

## Abstract

**Introduction:** Associations between breastfeeding and brain development, in the context of child, perinatal, and sociodemographic variables, remain unclear. This study investigates whether exclusive breastfeeding for the first 6 months and total duration of any breastfeeding are associated with brain white matter microstructure in young children.

**Methods:** This study included a sample of 83 mothers and 85 typically developing children (42 males). Children underwent their first diffusion tensor imaging scan between ages 2.34-6.97 years; some children returned multiple times, providing a total of 331 datasets. Feeding information was collected from the mothers at 3, 6, and 12 months postpartum and at their child’s scan to calculate breastfeeding status at 6 months (exclusive or not) as well as total duration of any breastfeeding. Linear regression was used to investigate associations between breastfeeding exclusivity/duration and fractional anisotropy (FA, a measure sensitive to myelination/axonal packing/fibre coherence) for the whole brain and 10 individual white matter tracts.

**Results:** Breastfeeding exclusivity and duration were associated with global and regional white matter microstructure, even after controlling for perinatal and sociodemographic factors. Greater exclusivity was associated with higher FA in females and lower FA in males.

**Conclusions:** These findings suggest white matter differences associated with breastfeeding that differ by sex. These may stem from different trajectories in white matter development between males and females in early childhood and suggest possible long-term white matter differences associated with breastfeeding.

## 1. Introduction

Breastfeeding is associated with positive health outcomes in children (Kramer, 2010) along with health and economic advantages for mothers (Martin et al., 2016; McInerny, 2014). Accordingly, the World Health Organization (WHO) recommends exclusive breastfeeding for the first 6 months of life followed by continued breastfeeding with appropriate complementary foods up to 2 years of age or beyond (WHO, 2003). There is some evidence that greater exclusivity in the first 6 months and longer breastfeeding are associated with cognitive advantages for children and adolescents (Anderson et al., 1999; Bernard et al., 2013; Kramer et al., 2008), possibly stemming from the nutritional composition of breastmilk and/or caregiver interactions during breastfeeding promoting cognitive and brain development (Guesnet et al., 2011; Krol et al., 2018).

Neuroimaging offers insight into the brain structures that have been associated with cognitive abilities; however, only a few magnetic resonance imaging (MRI) studies have probed the underlying associations between brain structure and breastfeeding. These studies highlight the role of white matter, suggesting that exclusivity and duration of breastfeeding are associated with alterations to myelin water fraction (Deoni et al., 2018; Deoni et al., 2013), anisotropy and diffusivity (Bauer et al., 2020; Blesa et al., 2019; Ou et al., 2014), and total white matter volume (Isaacs et al., 2010). Specifically, greater myelination of frontal-temporal white matter in full-term infants and children (Deoni et al., 2018; Deoni et al., 2013) and higher fractional anisotropy (FA; a measure associated with myelination, axonal packing, and fibre coherence) in frontal-parietal white matter of preterm infants (Blesa et al., 2019) and frontal-temporal white matter of school-aged children (Bauer et al., 2020) have all been associated with breastfeeding. Other studies have reported null findings: white matter volume did not differ between breastfed and non-breastfed full-term-born adolescents (Luby et al., 2016), nor was it associated with duration of breastfeeding in school-aged children (Bauer et al., 2020) or dosage of breast milk in preterm infants scanned at term-equivalent age (Belfort et al., 2016; Blesa et al., 2019) and at 7 years of age (Belfort et al., 2016). Furthermore, only some studies have examined sex differences; one showed that the percentage of expressed maternal breastmilk in preterm-born infants’ diets was associated with white matter volume in males but not females during adolescence (Isaacs et al., 2010), while the other showed that breastfed males had significantly higher whole brain FA than formula-fed males aged 7-8, with no group differences in females (Ou et al., 2014). Given that males and females show differing trajectories of brain development (Reynolds et al., 2019), it is critical to disentangle the potentially complex associations among white matter and breastfeeding, in the context of child age and sex, in large, longitudinal prospective studies.

It is also important to further investigate the role of perinatal and sociodemographic variables in breastfeeding-brain structure associations. Patterns of breastfeeding vary with maternal education and intelligence, child’s gestational age and birthweight, as well as socioeconomic status (Der et al., 2006; Jessri et al., 2013) and importantly, these variables are all associated with neurodevelopment (Batalle et al., 2017; Brito et al., 2014; Lugo-gil et al., 2008; Noble et al., 2015; Ronfani et al., 2015). Some studies that accounted for confounding variables such as maternal intelligence or education or household income observed attenuated associations between breastfeeding and cognitive abilities (Gibson-Davis et al., 2006; Horwood et al., 2001; Jacobson et al., 1999; McCrory et al., 2013). Thus, it is critical to control for perinatal and sociodemographic factors when investigating associations between neurodevelopment and breastfeeding in pediatric populations.

Of particular interest is understanding whether breastfeeding is associated with brain development during the early childhood period, a time of extensive brain changes and rapid cognitive, behavioral and emotional development (Deoni et al., 2012; Hermoye et al., 2006; Pfefferbaum et al., 1994). The objective of this study was to assess the association between white matter microstructure in young children (2-7 years) and breastfeeding exclusivity status at 6 months of age, as well as total duration of any breastfeeding. We used diffusion tensor imaging (DTI) to assess tissue microstructure as this technique is sensitive to myelination, axonal packing, and fibre coherence. Given the diverse and widespread nature of previous white matter alterations associated with breastfeeding (Bauer et al., 2020; Belfort et al., 2016; Blesa et al., 2019; Deoni et al., 2018; Deoni et al., 2013; Ou et al., 2014), we investigated microstructure of white matter tracts across the brain, as well as the whole brain average, accounting for age-related changes and sex differences in the context of perinatal and sociodemographic variables.

Based on the literature showing global and regional differences in myelin water fraction and white matter anisotropy (Bauer et al., 2020; Blesa et al., 2019; Deoni et al., 2018; Deoni et al., 2013; Ou et al., 2014), we hypothesized that exclusive breastfeeding for the first 6 months and longer durations of breastfeeding would be associated with higher FA in the whole brain and in individual white matter tracts.

## 2. Methods

### 2.1 Participants

This study included a convenience sample of 85 typically-developing children (42 males; 2 sets of non-twin siblings) and their 83 mothers. Participants were recruited from Calgary and surrounding areas (n=8) as well as from the ongoing Alberta Pregnancy Outcomes and Nutrition (APrON) study (n=77), a longitudinal cohort study of pregnant women (Kaplan et al., 2014). Inclusion criteria were full term birth (>37 weeks’ gestation), birthweight >2500 grams, living in the Calgary area, and ability to speak English as a primary language. Exclusion criteria were contraindications to MRI (e.g. braces, metal implants), brain injury, or a diagnosis of genetic or developmental disorders that impact cognitive and/or motor function (e.g. autism spectrum disorder). At the time of the first MRI scan, children were between 2.34 and 6.97 years (3.88±0.94 years); families were invited to return approximately every 6 months for a follow-up scan, though not everyone returned for each visit. This study included a total of 331 datasets from the 85 participants, with an average time of 7.52±4.43 months between visits. Data analyzed here includes 19 children with one scan, 15 with two scans, 4 with three scans, 11 with four scans, 16 with five scans, 11 with six scans, 7 with seven scans, 1 with eleven scans, and 1 with twenty scans. Written parental/guardian informed consent and verbal child assent were obtained for each subject at each scan. The University of Calgary Conjoint Health Research Ethics Board (CHREB) approved this study (REB18-0647).

### 2.2 Data Collection of Breastfeeding Patterns

For participants recruited from the APrON study, prospective data was acquired from the mothers about breastfeeding when children were 3 months, 6 months, and 12 months of age. Mothers were asked about breastfeeding and formula feeding patterns, including child’s age when they started and stopped, along with frequency of feeding, as outlined in Figure 1. The questionnaire also asked when and if other liquids (non-breast milk and non-milk liquids) or solid foods were introduced and their frequency of consumption. For all participants, mothers were asked again via a questionnaire about the child’s age when they started and stopped breastfeeding and formula feeding during the appointment for their child’s MRI scan.

**Figure 1.**
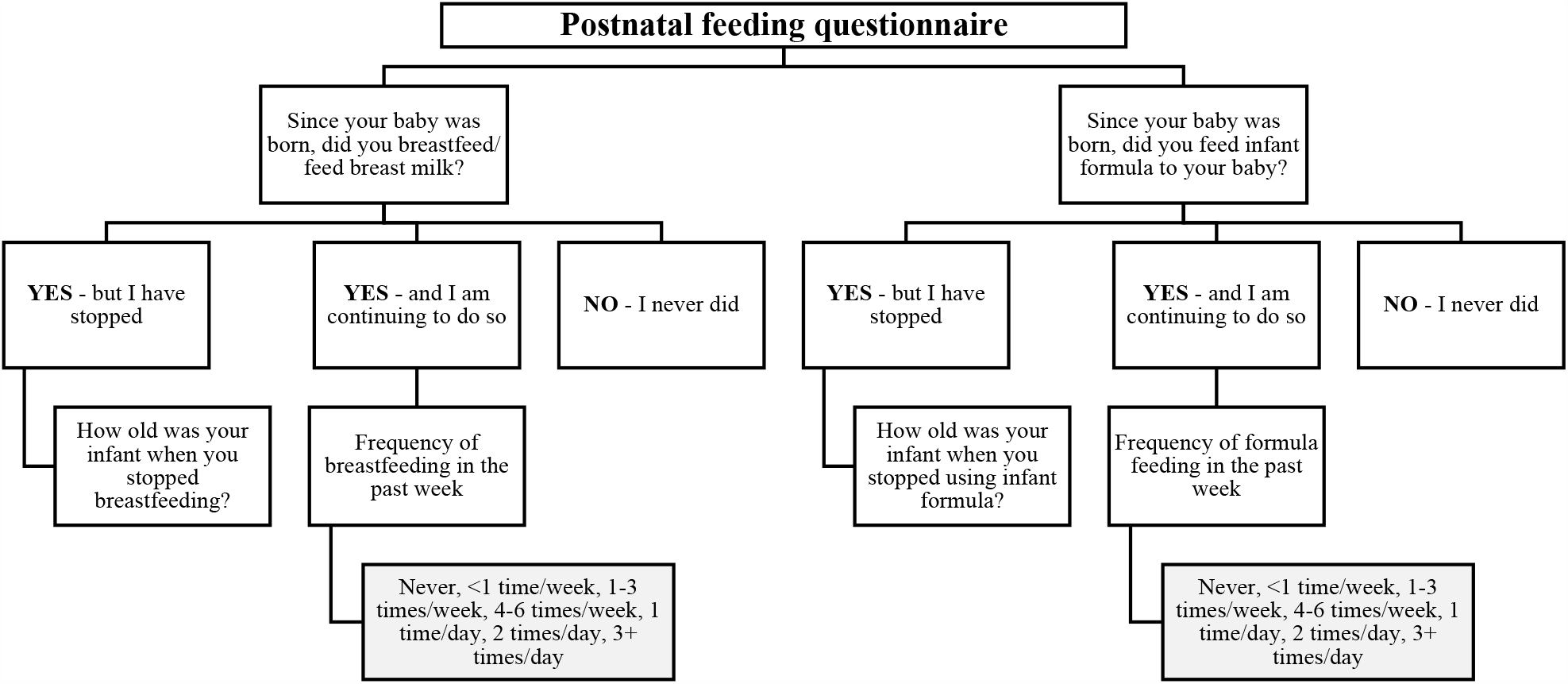
Infant feeding questionnaire included in the Alberta Pregnancy Outcomes and Nutrition (APrON) study. Participants were asked at 3, 6, and 12 months about feeding patterns for breastmilk and infant formula.

### 2.3 Data Collection of Child, Perinatal and Sociodemographic Covariates

At the first MRI scan, information about mothers (age at delivery, ethnicity) and children (age at scan, sex, gestational age at birth, birthweight) was obtained as a part of questionnaires. To assess socioeconomic status, information was collected about household income in the past year as a categorical variable (<$25,000; $25,000-49,999; $50,000-74,999;$75,000-99,999;$100,000-124,999; $125,000-149,999; $150,000-174,999; >$175,000) and maternal years of post-secondary education as a continuous variable. A subset of children (n=73) completed the Wechsler Preschool and Primary Scale of Intelligence–Fourth Edition (WPPSI-IV)(Wechsler, 2012) within 24 months of their MRI scan, which provided a full-scale intelligence quotient (FSIQ).

### 2.4 Neuroimaging Data Acquisition

MRI scanning took place at the Alberta Children’s Hospital on the research-dedicated GE 3T MR750w system with a 32-channel head coil. Children were not sedated for scanning, but were prepared using at-home materials and/or mock scanner training (Thieba et al., 2018). Whole-brain diffusion weighted images were acquired in 4:03 minutes using a single shot spin echo echo-planar imaging sequence with: 1.6 x 1.6 x 2.2 mm resolution (resampled to 0.78 x 0.78 x 2.2 mm on the scanner), TR = 6750 ms; TE = 79 ms, 30 gradient encoding directions at b=750 s/mm^2^, and five interleaved images without diffusion encoding at b=0 s/mm^2^. Diffusion MRI was acquired as part of a longer neuroimaging protocol (Reynolds et al., 2020).

### 2.5 Neuroimaging Data Processing

DTI data was visually quality checked, and volumes with artifacts or motion corruption were removed, as per our previous methods.(Reynolds et al., 2019; Walton et al., 2018). All children had ≥18 diffusion weighted volumes (28±3) and ≥2 non-diffusion weighted volumes (5±0) remaining after removal of motion-corrupted data. Data was preprocessed using ExploreDTI (V4.8.6), including corrections for signal drift, Gibbs ringing, subject motion, and eddy current distortions (Leemans et al., 2009). Semi-automated deterministic streamline tractography was used to delineate 10 major white matter tracts: the corpus callosum (genu, body, splenium), fornix, cingulum, pyramidal tract, superior and inferior longitudinal fasciculi (SLF, ILF), inferior fronto-occipital fasciculus (IFOF), and uncinate fasciculus (UF) (Lebel et al., 2012; Lebel et al., 2008; Reynolds et al., 2019). Detailed descriptions of region of interest selection for tracking procedures can be found at https://doi.org/10.6084/m9.figshare.7603271 (Reynolds et al., 2020). The minimum FA threshold was set to 0.20 to initiate and continue tracking, and angle threshold set to 30° to minimize spurious fibers (Lebel et al., 2008; Reynolds et al., 2019). Tracts were manually quality checked and additional exclusion regions of interest were drawn as required. Average FA was extracted for all white matter fibers in the whole brain as well as for each of the 10 major tracts. For every tract except the corpus callosum (genu, body, splenium) and the fornix, FA values were calculated separately for left and right hemispheres and subsequently averaged. As additional measures of white matter microstructure, mean, axial and radial diffusivity (MD, AD, RD) were also extracted for all white matter tracts (see supplementary Methods).

### 2.6 Statistical Analysis

All statistical analyses were completed using Statistical Package for the Social Sciences (SPSS; Version 25).

#### 2.6.1 Demographics

The Shapiro-Wilk test was used to test all variables for normality, which showed that the following variables were not normally distributed: age at scan, gestational age at birth, age at delivery, post-secondary education, total duration of breastfeeding. Thus, we used a Mann-Whitney U test to compare these variables between breastfeeding exclusivity groups (described in next section). For the remaining variables (birthweight, FSIQ) that were normally distributed, we used a t-test when comparing between breastfeeding exclusivity groups. Chi-squared tests were used to test for group differences for child’s sex, maternal ethnicity, and family income. The Shapiro-Wilk test determined that total duration of breastfeeding was not normally distributed. Thus, Spearman correlations were used to test for associations between total duration of any breastfeeding with child’s age at scan, gestational age, birthweight, FSIQ, maternal years of post-secondary education and maternal age at delivery. A univariate test of variance was used to test for associations between total duration of any breastfeeding and child’s sex, maternal ethnicity, and family income. These analyses were completed for the entire sample (n=85) and for the subset of participants with FSIQ scores (n=73).

#### 2.6.2 Breastfeeding exclusivity analysis

Children were divided into groups based on the information provided by mothers about feeding patterns at the 3-months and 6-months of age. Exclusive breastfeeding (EBF) was defined as feeding breastmilk with no non-breastmilk milk liquids (e.g., formula, cow’s milk) for the first 6 months of life; this group consisted of 50 children (200 datasets). Children who were not exclusively breastfed (nEBF) for the first 6 months of life received either a mix of breastmilk and formula (n=29; 108 datasets) or were predominantly formula-fed (n=6; 23 datasets). A general linear mixed effects analysis was used to test group differences (EBF vs. nEBF) for FA of the whole brain and individual tracts. For this analysis, the primary model included breastfeeding exclusivity group, child’s sex, child’s age at scan, as well as all possible group-sex-age interactions (sex∗age, group∗sex, group∗age, group∗sex∗age); subject was included as a random factor. Interactions that were not significant were removed and the model was re-run. Where sex interactions were significant, post-hoc tests were run separately in males and in females to better understand sex differences in the association between breastfeeding exclusivity group and FA. Significance level was set at p<0.05; false discovery rate was used to correct for 22 multiple comparisons (2 analyses – exclusivity and duration – for each of the 10 tracts and the whole brain). Identical analyses were conducted with MD, AD and RD (see supplementary Methods).

#### 2.6.3 Breastfeeding duration analysis

Total duration of any breastfeeding was available for 82 children (327 datasets) and ranged from 0.008 years to 4 years (1.2±0.7 years; all children were breastfed at least once). A general linear mixed effects model was used to test associations between total duration of any breastfeeding and FA in the whole brain and individual tracts. This analysis tested breastfeeding as a continuous variable, with the primary model including duration of breastfeeding, child’s sex, child’s age at scan, as well as all duration-sex-age interactions (sex∗age, duration∗sex, duration∗age, and duration∗sex∗age); subject was included as a random factor. Interactions that were not significant were removed and the model was re-run. Significance level was set at p<0.05; false discovery rate was used to correct for 22 multiple comparisons (2 analyses – exclusivity and duration – for each of the 10 tracts and the whole brain). Identical analyses were conducted with MD, AD and RD (see supplementary Methods).

#### 2.6.4 Supplementary analyses

Follow-up analyses were run for both breastfeeding exclusivity group and breastfeeding duration analyses, adding the following variables to the primary statistical model: child’s gestational age, child’s birthweight, maternal years of post-secondary education, maternal age at delivery, maternal ethnicity, and family income. Covariates were included based on known associations with patterns of breastfeeding (Der et al., 2006; Gibbs et al., 2014; Jain et al., 2002; Jessri et al., 2013; Walfisch et al., 2013) and/or neurodevelopmental outcomes (Batalle et al., 2017; Brito et al., 2014; Lugo-gil et al., 2008; Noble et al., 2015; Ronfani et al., 2015). Two additional analyses were also run; the first, adding child’s FSIQ to the primary statistical model for those children who had FSIQ scores (n=73; 296 datasets) and the second, adding the number of diffusion weighted volumes remaining for each dataset to the primary statistical model to account for motion.

### 2.7 Data availability

Neuroimaging data is publicly available on the Open Science Framework here: https://osf.io/axz5r/ (Reynolds et al., 2020). Alberta Pregnant Outcomes and Nutrition (APrON) cohort breastfeeding data is available on SAGE: https://dataverse.library.ualberta.ca/dataverse/SAGE.

## 3. Results

### 3.1 Demographics

The EBF group had more females (p=0.003), higher gestational age at birth (p=0.028), and were breastfed for longer durations (p<0.001), compared to the nEBF group (Table 1). Group differences and correlations between breastfeeding and demographic variables within the subset of participants with FSIQ scores showed similar results to the entire sample; however, gestational age at birth was no longer significantly different between groups.

**Table 1.**
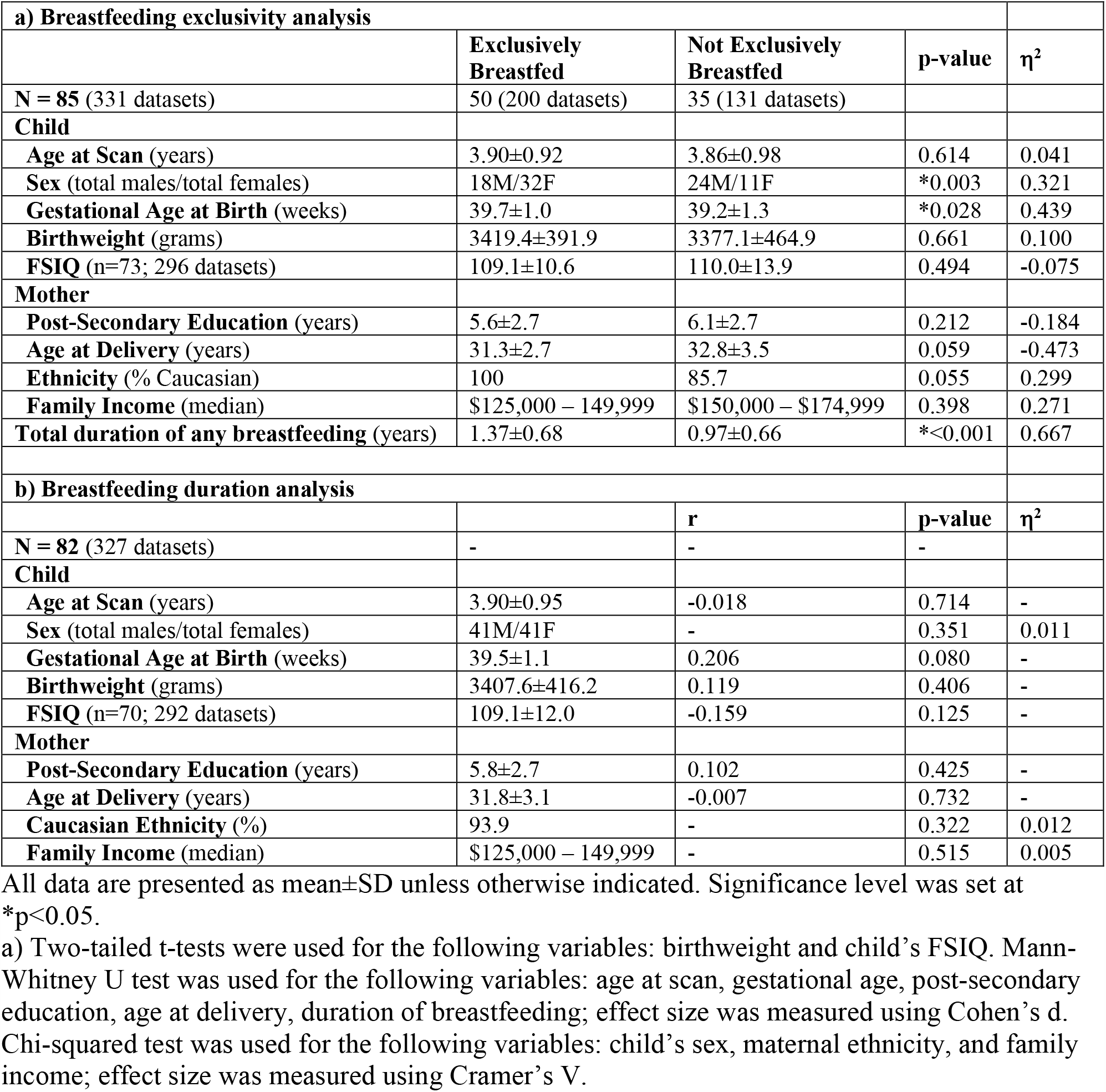
Participant Characteristics.

| **a) Breastfeeding exclusivity analysis** |  |  |  |  |
| --- | --- | --- | --- | --- |
|  | **Exclusively Breastfed** | **Not Exclusively Breastfed** | **p-value** | **$\eta^2$** |
| **N = 85** (331 datasets) | 50 (200 datasets) | 35 (131 datasets) |  |  |
| **Child** |  |  |  |  |
| **Age at Scan** (years) | 3.90±0.92 | 3.86±0.98 | 0.614 | 0.041 |
| **Sex** (total males/total females) | 18M/32F | 24M/11F | *0.003 | 0.321 |
| **Gestational Age at Birth** (weeks) | 39.7±1.0 | 39.2±1.3 | *0.028 | 0.439 |
| **Birthweight** (grams) | 3419.4±391.9 | 3377.1±464.9 | 0.661 | 0.100 |
| **FSIQ** (n=73; 296 datasets) | 109.1±10.6 | 110.0±13.9 | 0.494 | -0.075 |
| **Mother** |  |  |  |  |
| **Post-Secondary Education** (years) | 5.6±2.7 | 6.1±2.7 | 0.212 | -0.184 |
| **Age at Delivery** (years) | 31.3±2.7 | 32.8±3.5 | 0.059 | -0.473 |
| **Ethnicity** (% Caucasian) | 100 | 85.7 | 0.055 | 0.299 |
| **Family Income** (median) | \$125,000 – 149,999 | \$150,000 – \$174,999 | 0.398 | 0.271 |
| **Total duration of any breastfeeding** (years) | 1.37±0.68 | 0.97±0.66 | *<0.001 | 0.667 |
| **b) Breastfeeding duration analysis** |  |  |  |  |
|  |  | **r** | **p-value** | **$\eta^2$** |
| **N = 82** (327 datasets) | - | - | - |  |
| **Child** |  |  |  |  |
| **Age at Scan** (years) | 3.90±0.95 | -0.018 | 0.714 | - |
| **Sex** (total males/total females) | 41M/41F | - | 0.351 | 0.011 |
| **Gestational Age at Birth** (weeks) | 39.5±1.1 | 0.206 | 0.080 | - |
| **Birthweight** (grams) | 3407.6±416.2 | 0.119 | 0.406 | - |
| **FSIQ** (n=70; 292 datasets) | 109.1±12.0 | -0.159 | 0.125 | - |
| **Mother** |  |  |  |  |
| **Post-Secondary Education** (years) | 5.8±2.7 | 0.102 | 0.425 | - |
| **Age at Delivery** (years) | 31.8±3.1 | -0.007 | 0.732 | - |
| **Caucasian Ethnicity** (%) | 93.9 | - | 0.322 | 0.012 |
| **Family Income** (median) | \$125,000 – 149,999 | - | 0.515 | 0.005 |

All data are presented as mean ± SD unless otherwise indicated. Significance level was set at \* $p<0.05$ .
a) Two-tailed t-tests were used for the following variables: birthweight and child's FSIQ. Mann-Whitney U test was used for the following variables: age at scan, gestational age, post-secondary education, age at delivery, duration of breastfeeding; effect size was measured using Cohen's d. Chi-squared test was used for the following variables: child's sex, maternal ethnicity, and family income; effect size was measured using Cramer's V.

### 3.2 Breastfeeding exclusivity analysis

Group-sex interactions were significant for whole brain FA (p<0.001; Table 2). Specifically, nEBF males had significantly higher FA than EBF males, whereas EBF females had higher FA than nEBF females (Figure 2). A similar group-sex interaction was also significant for FA in most tracts (Table 2), with EBF females > nEBF females and EBF males < nEBF males (Figure 2). Exceptions occurred in the fornix, the UF and the SLF, where only females showed significant group differences for FA, and the ILF, where only males showed significant group differences for FA. The group-age interaction was significant in the body of the corpus callosum (p=0.046), the pyramidal tract (p<0.001), and the IFOF (p=0.009); the latter two tracts show steeper increases in FA with age in the EBF group (Figure 3). All two-way interactions survived correction for multiple comparisons with the exception of the group-age interaction in the body of the corpus callosum. All findings remained significant after controlling for perinatal/sociodemographic covariates, child’s FSIQ, and motion (number of volumes removed) with the exception of the group-age interaction in the IFOF, which did not remain significant after controlling for child’s FSIQ. Results for MD, AD and RD are shown in the supplementary material (Results, Table S1, Table S2, Table S3, Figure S1 and Figure S2).

**Table 2.** Linear mixed effects results analyzing differences in FA between breastfeeding groups (i.e. EBF vs. nEBF).

|  | Fractional Anisotropy |  |  |  |  |
| --- | --- | --- | --- | --- | --- |
|  | Main Effects |  |  | Interactions |  |
| **Whole Brain** | **Age** | **Sex** | **Group** | **Group\*Sex** | **Group\*Age** |
| Parameter Estimate | 6.0E-03 | 8.6E-03 | 9.1E-03 | -1.5E-02 | - |
| t | 16.117 | 4.188 | 4.760 | -6.803 | - |
| 95% CI ($10^{-3}$) | 5.3, 6.7 | 4.6, 12.6 | 5.3, 12.8 | -20.0, -11.0 | - |
| p-value | <0.001 | 0.459 | 0.247 | <0.001 | - |
| **CC Genu** |  |  |  |  |  |
| Parameter Estimate | 8.4E-03 | 2.3E-02 | 1.2E-02 | -2.2E-02 | - |
| t | 10.834 | 6.004 | 3.244 | -4.986 | - |
| 95% CI ($10^{-3}$) | 6.7, 10.0 | 15.4, 30.5 | 4.6, 18.9 | -30.8, -13.2 | - |
| p-value | <0.001 | <0.001 | 0.734 | <0.001 | - |
| **CC Body** |  |  |  |  |  |
| Parameter Estimate | 4.6E-03 | -3.7E-03 | 9.8E-03 | -2.1E-02 | 3.4E-03 |
| t | 3.561 | -0.422 | 2.574 | -4.542 | 2.008 |
| 95% CI ($10^{-3}$) | 2.1, 7.1 | -21.2, 13.7 | 2.3, 17.4 | -30.2, -11.9 | 0.06, 6.7 |
| p-value | <0.001 | 0.085 | 0.767 | <0.001 | 0.046 |
| **CC Splenium** |  |  |  |  |  |
| Parameter Estimate | 5.4E-03 | 8.6E-03 | 6.7E-03 | -1.2E-02 | - |
| t | 17.045 | 2.502 | 2.199 | -3.564 | - |
| 95% CI ($10^{-3}$) | -72.2, 72.2 | 1.8, 15.3 | 0.7, 12.7 | 19.6, -5.3 | - |
| p-value | 0.688 | 0.188 | 0.785 | 0.001 | - |
| **Fornix** |  |  |  |  |  |
| Parameter Estimate | 7.6E-03 | 1.0E-02 | 1.4E-02 | -1.6E-02 | - |
| t | 8.497 | 2.498 | 3.404 | -3.261 | - |
| 95% CI ($10^{-3}$) | 5.8, 9.4 | 21.9, 18.5 | 5.8, 21.8 | -26.0, -6.4 | - |
| p-value | <0.001 | 0.372 | 0.022 | 0.001 | - |
| **Cingulum** |  |  |  |  |  |
| Parameter Estimate | 8.3E-03 | 1.9E-02 | 1.9E-02 | -2.7E-02 | - |
| t | 9.036 | 4.502 | 4.596 | -5.213 | - |
| 95% CI ($10^{-3}$) | 6.5, 10.1 | 10.9, 27.8 | 11.1, 27.8 | -37.1, -16.8 | - |
| p-value | <0.001 | 0.025 | 0.021 | <0.001 | - |
| **IFOF** |  |  |  |  |  |
| Parameter Estimate | 6.5E-03 | 5.8E-03 | -2.0E-03 | -2.0E-02 | 3.2E-03 |
| t | 6.587 | 1.909 | -0.313 | -5.727 | 2.645 |
| 95% CI ($10^{-3}$) | 4.6, 8.5 | -0.2, 11.7 | -14.4, 10.5 | -26.7, -13.0 | 0.8, 5.6 |
| p-value | <0.001 | 0.017 | 0.047 | <0.001 | 0.009 |
| **ILF** |  |  |  |  |  |
| Parameter Estimate | 1.1E-02 | 4.7E-04 | 6.0E-03 | -1.3E-02 | - |
| t | 15.739 | 0.137 | 1.909 | -3.324 | - |
| 95% CI ($10^{-3}$) | 9.7, 12.5 | -6.2, 7.2 | -0.2, 12.1 | -20.8, -5.3 | - |
| p-value | <0.001 | 0.002 | 0.788 | 0.001 | - |

|  | Fractional Anisotropy (continued) |  |  |  |  |
| --- | --- | --- | --- | --- | --- |
|  | Main Effects |  |  | Interactions |  |
|  | Age | Sex | Group | Group*Sex | Group*Age |
| **SLF** |  |  |  |  |  |
| Parameter Estimate | 6.2E-03 | 1.1E-03 | 1.2E-02 | -1.4E-02 | - |
| t | 9.267 | 0.322 | 3.596 | -3.454 | - |
| 95% CI ($10^{-3}$) | 4.9, 7.5 | -5.5, 7.7 | 5.4, 18.4 | -21.5, -5.9 | - |
| p-value | <0.001 | 0.004 | 0.012 | <0.001 | - |
| **Pyramidal Tract** |  |  |  |  |  |
| Parameter Estimate | 4.8E-03 | 1.4E-02 | -1.3E-02 | -2.0E-02 | 5.1E-03 |
| t | 4.912 | 5.071 | -2.071 | -6.223 | 4.299 |
| 95% CI ($10^{-3}$) | 2.9, 6.8 | 8.7, 19.8 | -24.8, -0.6 | -26.2, -13.6 | 2.7, 7.4 |
| p-value | <0.001 | 0.008 | <0.001 | <0.001 | <0.001 |
| **UF** |  |  |  |  |  |
| Parameter Estimate | 6.1E-03 | 8.3E-03 | 1.8E-02 | -1.6E-02 | - |
| t | 9.196 | 2.522 | 5.585 | -4.326 | - |
| 95% CI ($10^{-3}$) | 4.8, 7.4 | 1.8, 14.8 | 11.5, 24.0 | -24.0, -9.0 | - |
| p-value | <0.001 | 0.977 | <0.001 | <0.001 | - |

If interactions were not significant, they were removed from the model.

**Figure 2.**
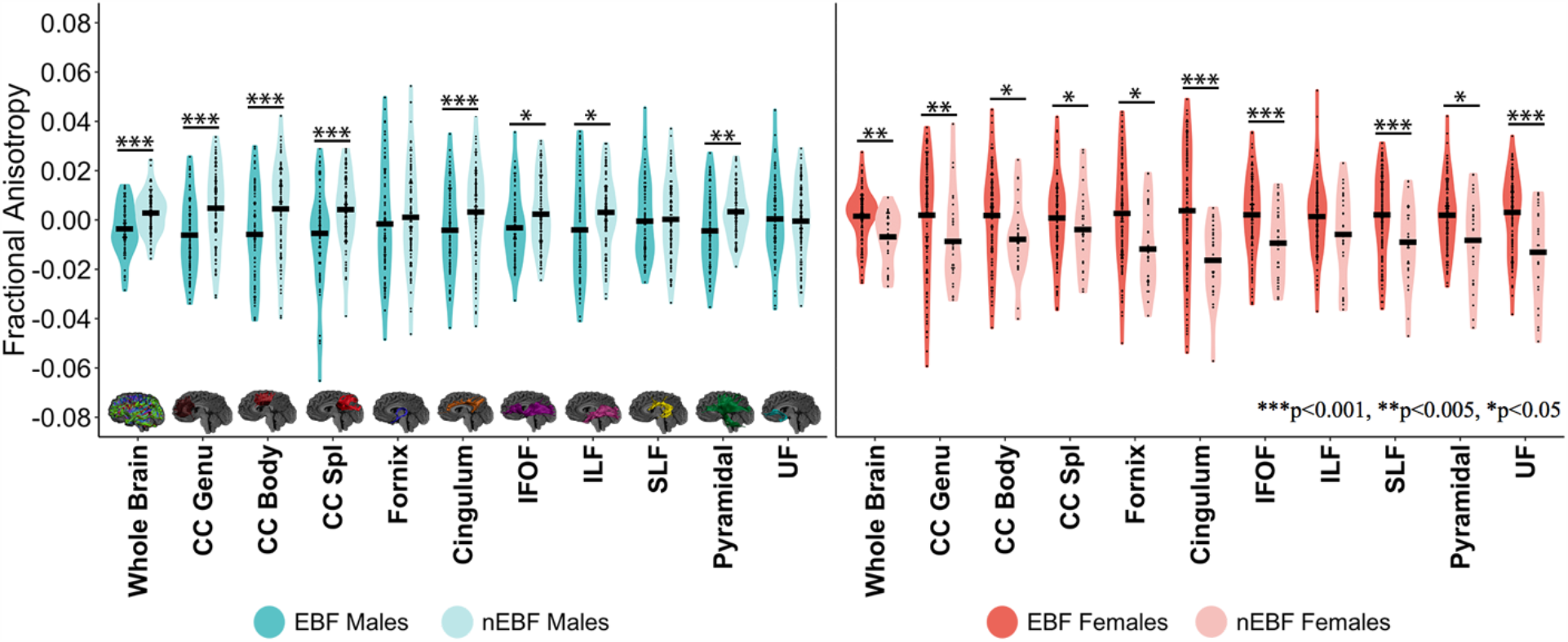
Breastfeeding group and sex interactions for fractional anisotropy in the whole brain and individual white matter tracts. EBF males showed significantly lower FA than nEBF males in the whole brain and all individual tracts except the fornix, the SLF and the UF. EBF females showed significantly higher FA than nEBF females in the whole brain and all individual tracts except the ILF. Data is represented as mean ± 95% confidence interval for FA values for whole brain and individual tracts in EBF males vs. nEBF males and EBF females vs. nEBF females. FA values are corrected for age at the participant’s scan. CC Genu = genu of the corpus callosum, CC Body = body of the corpus callosum, CC Splenium = splenium of the corpus callosum. ∗∗∗p<0.001; ∗∗p<0.005; ∗p<0.05.

**Figure 3.**
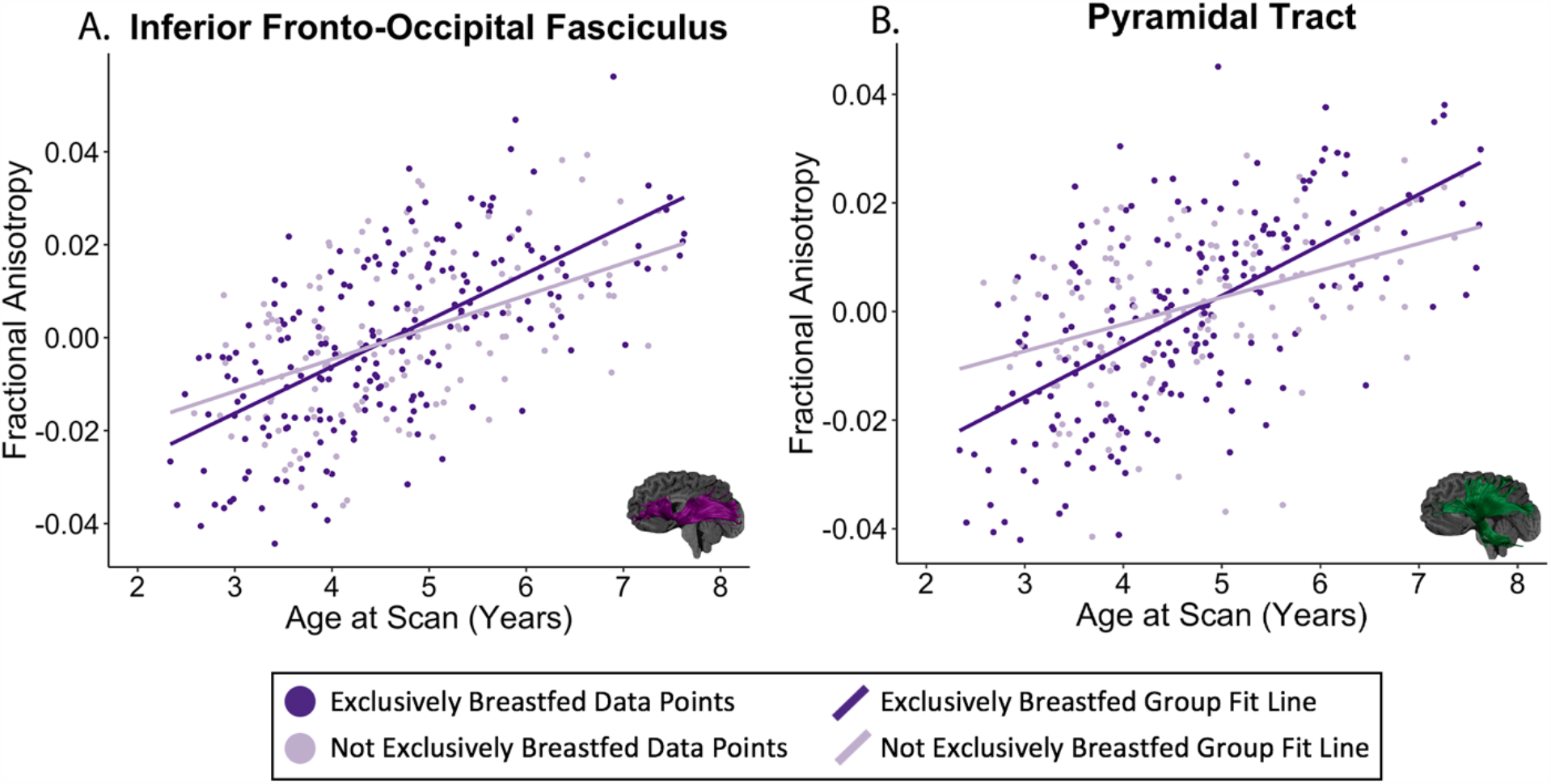
Significant interaction between breastfeeding group and age for FA in the IFOF, F=7.0, p=0.009 (a) and the pyramidal tract, F=18.5, p<0.001 (b). In both tracts, the exclusively breastfed group shows steeper increases in FA with age. All FA values are corrected for sex.

### 3.3 Breastfeeding duration analysis

The duration-sex interaction was significant for FA in the body of the corpus callosum (p=0.009), the cingulum (p=0.015) and the ILF (p=0.004; Table 3). As well, the duration-age interaction was significant for FA in the pyramidal tract (p=0.006; Table 3) and there was a main effect of duration for FA in the SLF (p=0.017; Table 3). The only finding that survived correction of multiple comparisons was the ILF, showing that lower FA was associated with longer breastfeeding for females, while no associations were noted between FA and duration with males (Figure 4). This finding remained significant after controlling for perinatal/sociodemographic covariates, child’s FSIQ, and motion (number of volumes removed). Additionally, as a small number of participants (6 participants; 16 datasets) were breastfed considerably longer than average, we re-ran the analysis without those participants; all results remained significant. MD, AD and RD results are shown in the supplementary material (Results, Table S4, Table S5, Table S6 and Figure S3).

**Table 3.**
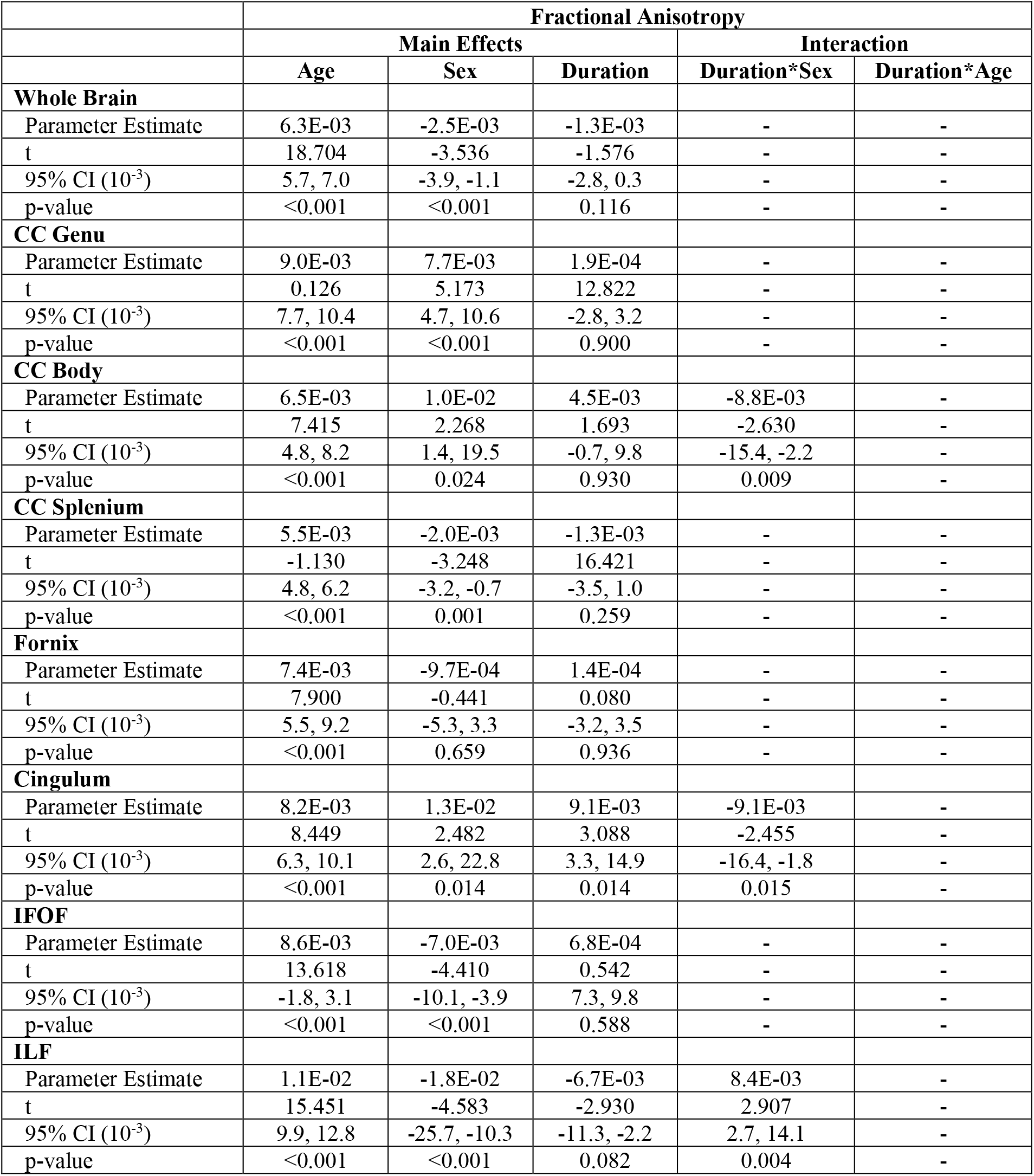
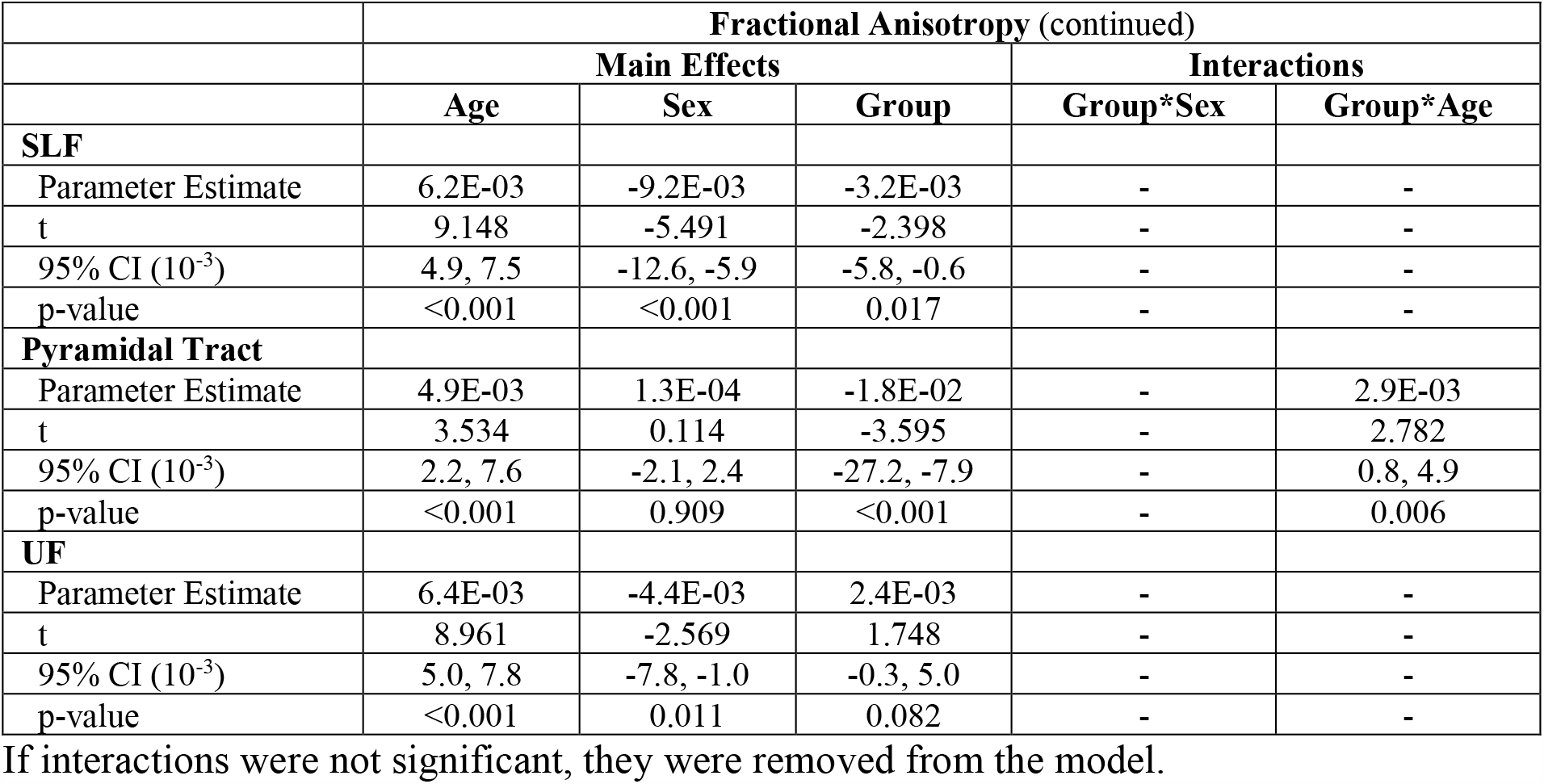
Linear mixed effects results analyzing associations between FA and total duration of any breastfeeding.

**Figure 4.**
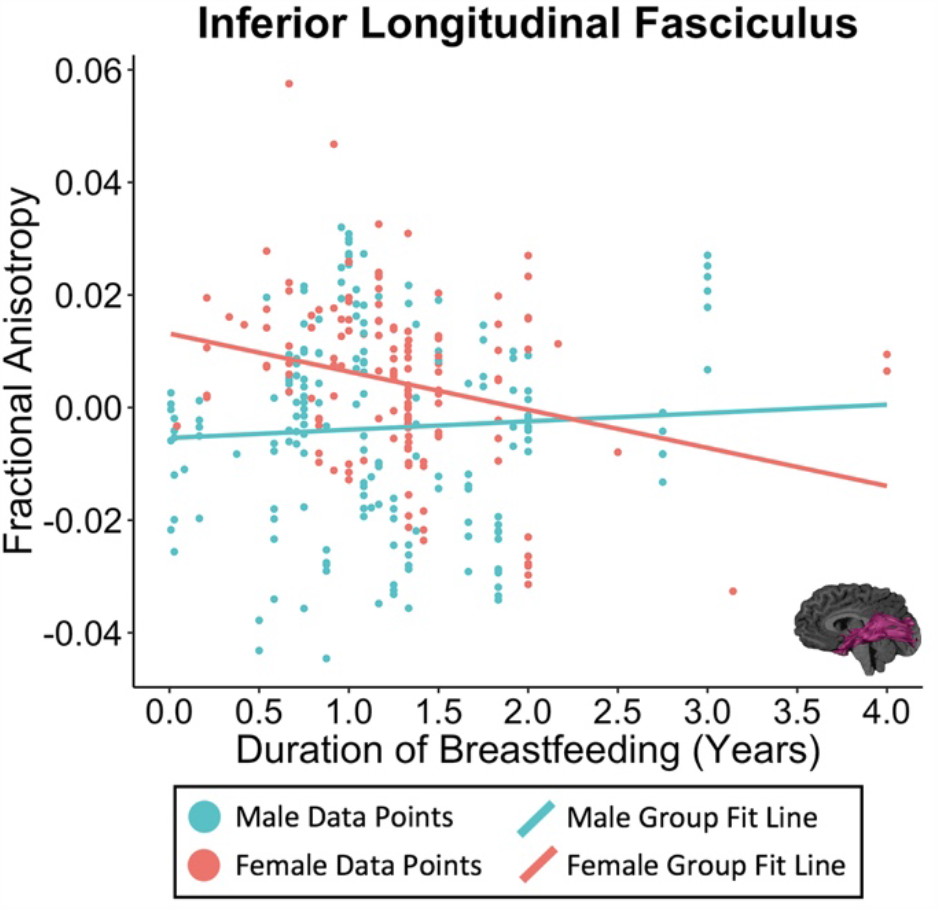
Significant interactions between breastfeeding duration and child’s sex for FA in the inferior longitudinal fasciculus (F=8.5, p=0.004), where FA decreases as duration of breastfeeding increases for females. All FA values are corrected for age at the participant’s scan.

## 4. Discussion

In this prospective longitudinal study, exclusive breastfeeding for the first 6 months of life and longer durations of breastfeeding were associated with altered white matter structure in early childhood. Most findings remained significant after accounting for potential confounders and revealed that exclusive breastfeeding is associated with sex-specific white matter differences in young children.

Females who were EBF for the first 6 months had significantly higher FA on both a global and regional level compared to nEBF females. The opposite was observed in males: nEBF boys had significantly higher FA at a global and regional level. Our findings in females align with previous research in full-term and preterm infants and young children where breastfeeding exclusivity and/or duration was associated with higher myelin water fraction (Deoni et al., 2018; Deoni et al., 2013) or higher FA (Bauer et al., 2020; Blesa et al., 2019). Interestingly, in studies that examined sex differences, no associations were reported between breastfeeding and white matter microstructure in term-born school-aged females (Ou et al., 2014) or white matter volume in preterm adolescent females (Isaacs et al., 2010). For males, our results are inconsistent with previous research that reported associations between breastfeeding exclusivity and/or duration, and higher myelin water fraction (Deoni et al., 2018; Deoni et al., 2013), higher FA (Bauer et al., 2020; Blesa et al., 2019; Ou et al., 2014) or higher white matter volume (Isaacs et al., 2010).

Sex differences in white matter development may help explain the differing associations in females versus males. Throughout infancy, childhood and into early adulthood, FA increases throughout the brain (Dubois et al., 2014; Hermoye et al., 2006; Krogsrud et al., 2016; Lebel et al., 2008; Reynolds et al., 2019). In young children, these developmental processes vary slightly between sexes, with young boys showing faster rates of change (steeper slopes) than girls. On the contrary, young girls have higher initial maturity (i.e. higher FA), suggesting that they undergo earlier brain development compared to boys (Reynolds et al., 2019). Similar observations have been made in older children and adolescents (Clayden et al., 2012; Seunarine et al., 2015; Wang et al., 2012). It is possible that females are further along in terms of white matter maturation compared to males, which may lead to sexually dimorphic breastfeeding-brain structure associations. As white matter continues to develop through late childhood and adolescence, it is possible that the association between breastfeeding and anisotropy will shift, such that older EBF males may have higher FA than older nEBF males. This would align with previous research findings in older, school-aged children (4- to 8.5-years-old), which noted that breastfed males had higher FA than formula-fed males in whole brain and regional white matter (Bauer et al., 2020; Ou et al., 2014). This is, in part, supported by the fact that the EBF group in our sample show steeper age-related increases in FA in some white matter tracts when compared to the nEBF group. Continued follow-up in this sample and more longitudinal studies will help to further understand associations between developmental trajectories of white matter and breastfeeding, clarifying whether the observed associations in young children persist across ages or whether they are unique to early childhood.

The most prominent differences between breastfeeding groups were in the cingulum, a tract associated with social-emotional processing and memory (Bubb et al., 2018) followed by the corpus callosum (Aboitiz et al., 2003), which is involved in integrating and transferring information between hemispheres. Other tracts with notable differences included the IFOF, which underlies visual processing and early language skills (Almairac et al., 2015), the pyramidal tract which underlies motor performance (Jang, 2014; Yeo et al., 2014), as well as fronto-temporal tracts, the SLF and the UF, which are associated with higher order cognitive functions, executive function, language, and social-emotional processing (Olson et al., 2015; Urger et al., 2015). Across numerous studies, breastfeeding has been associated with functions such as social-emotional processing (Oddy et al., 2010), motor performance (Belfort et al., 2016; Michels et al., 2017), cognition and memory (Belfort et al., 2016), as well as language (Dee et al., 2006), which may, in part, be associated with alterations to underlying white matter tracts that correspond to these functions.

One hypothesized mechanism by which breastfeeding may be associated with white matter development is the nutritional content of breastmilk. Breastmilk contains a variety of proteins, lipids, and other macro- and micro-nutrients that are necessary for white matter development (Chang et al., 2009; Hadley et al., 2016). For example, choline is abundant in breastmilk and is a precursor to phosphatidylcholine and sphingomyelin, which are required for the synthesis of the myelin sheath (Guesnet et al., 2011; Hoffman et al., 2009). Similarly, long-chain polyunsaturated fatty acids in breastmilk, specifically docosahexaenoic acid and arachidonic acid, support neuronal growth and myelination (Guesnet et al., 2011; Hoffman et al., 2009). It is possible that males and females respond differently to their early nutritional environments, which may subsequently alter neurodevelopment (Galante et al., 2018). Another possible mechanism is the increased opportunity for didactic and affectionate mother-child interactions during breastfeeding, which may contribute to the development of white matter tracts (Krol et al., 2018).

While these sex-specific findings were robust, there are limitations that are important to consider. Participants were primarily of moderate-to-high socioeconomic status and most identified as Caucasian; therefore, these results may not be representative of demographically different populations. While this sample limits the generalizability of our findings, it does ensure limited differences in perinatal and sociodemographic characteristics between the EBF and Nebf groups, allowing for better control for potential confounding factors. Furthermore, the feeding questions did not gather specific quantities of milk the child ingested from breastfeeding or formula-feeding, nor did they ask about method of feeding breastmilk (i.e., at the breast or via bottle), which could affect the associations observed here. This study also had few participants that were exclusively formula-fed, which may have reduced the power to detect group differences. Lastly, it is important to note that the duration of breastfeeding in our sample (14.3±8.4 months) is slightly, but not dramatically higher than durations reported in other neuroimaging studies with samples in the United States with 9.9±7.1 months (Bauer et al.,2020) or 12.5 ± 6.2 months (Ou et al., 2014) on average. These durations are all lower than the 2-year duration recommended by the WHO (WHO, 2003).

## 5. Conclusions

This study demonstrates that breastfeeding exclusivity during the first 6 months and total duration of breastfeeding are associated with sex-specific global and regional alterations to white matter development in young children. These complex associations should compel future research to comprehensively consider sex differences, age-related changes, and other covariates in their analysis. Understanding associations between brain development and breastfeeding is essential for health education and promotion strategies as these results suggest that breastfeeding is associated with long-term neurodevelopmental changes that persist into early childhood.

## Acknowledgements

The authors acknowledge the contributions of the APrON Study Team. We are extremely grateful to all the families who took part in this study and the whole APrON team (http://www.apronstudy.ca), investigators, research assistants, graduate and undergraduate students, volunteers, clerical staff and mangers. This cohort was established by an interdisciplinary team grant from Alberta Innovates Health Solutions (formally the Alberta Heritage Foundation for Medical Research) and additional funding (see above) assisted with the collection and analysis of data presented in this manuscript.

## Abbreviations

MRI: magnetic resonance imaging
FA: fractional anisotropy
DTI: diffusion tensor imaging
APrON: Alberta Pregnancy Outcomes and Nutrition
FSIQ: full-scale intelligence quotient
SLF: superior longitudinal fasciculi
ILF: inferior longitudinal fasciculus
IFOF: inferior fronto-occipital fasciculus
UF: and uncinate fasciculus
MD: mean diffusivity
EBF: exclusively breastfed
nEBF: not exclusively breastfed

