## Supplementary Data for "Association between breastfeeding during infancy and white matter microstructure in early childhood"

**Supplementary Material**

**S2. Methods**

As additional measures of white matter microstructure besides fractional anisotropy, mean, axial and radial diffusivity (MD, AD and RD) were also analyzed. Using ExploreDTI, average MD was extracted for all the white matter fibers in the whole brain as well as for each of the 10 major white matter tracts. Similarly, AD and RD were calculated for the 10 major white matter tracts. For every tract except the corpus callosum (body, splenium genu) and the fornix, MD, AD and RD values were calculated separately for left and right hemispheres and subsequently averaged. For all analyses, MD, AD and RD values were scaled by 1000 to bring them to a similar scale as fractional anisotropy (FA).

**S3. Results**

*S3.1 Breastfeeding exclusivity analysis*

MD: Group-sex interactions were significant for whole brain MD ($p<0.001$; Supplementary Table S3). Specifically, nEBF males had significantly lower MD than EBF males, whereas EBF females had lower MD than nEBF females (Supplementary Figure S1). The group-sex interaction was also significant for MD in most tracts (Supplementary Table S3), with EBF females > nEBF females and EBF males < nEBF males (Supplementary Figure S1). Exceptions occurred in the genu of the corpus callosum and the UF, where only females showed significant group differences for MD, as well as the whole brain, the body of the corpus callosum, the fornix and the pyramidal tract, where only males showed significant group differences for MD. The group-age interaction was significant in the genu of the corpus callosum ($p=0.007$), where EBF children showed steeper decreases in MD with age (Supplementary Figure S2A). The group-sex-age interaction was significant for MD of the fornix (p<0.001) and the pyramidal tract (p=0.016; (Supplementary Figure S2B, S2C). The majority of two- and three-way interactions survived correction for multiple comparisons and remained significant after controlling for perinatal/sociodemographic covariates and child's FSIQ. The only exception was the three-way interaction in the pyramidal tract that did not remain significant after controlling for perinatal/sociodemographic factors. The interactions in the genu and splenium of the corpus callosum, the UF and the SLF did not survive correction for motion (number of volumes removed).

**Supplementary Table S1. Linear mixed effects results analyzing differences in MD between breastfeeding groups (i.e. EBF vs. nEBF).**

| | **Mean Diffusivity (MD)** | | | | | |
|---|---|---|---|---|---|---|
| | **Main Effects** | | | **Interactions** | | |
| | **Age** | **Sex** | **Group** | **Group*Sex** | **Group*Age** | **Group*Sex*Age** |
| **Whole Brain** | | | | | | |
| Parameter Estimate | -1.3E-02 | -1.9E-03 | -1.3E-02 | 2.5E-02 | - | - |
| t | -3.370 | -0.470 | -17.598 | 5.657 | - | - |
| 95% CI ($10^{-3}$) | -20.2, -5.3 | -9.8, 6.0 | -14.8, -11.8 | 16.6, 34.3 | - | - |
| p-value | <0.001 | 0.639 | <0.001 | <0.001 | - | - |
| **CC Genu** | | | | | | |
| Parameter Estimate | -1.7E-02 | -3.9E-02 | 7.7E-04 | 5.0E-02 | -8.3E-03 | - |
| t | -6.593 | -4.959 | 0.047 | 5.827 | -2.749 | - |
| 95% CI ($10^{-3}$) | -21.6, -11.6 | -53.8, -23.2 | -31.3, 32.9 | 33.3, 67.3 | -14.3, -2.3 | - |
| p-value | <0.001 | <0.001 | 0.962 | <0.001 | 0.007 | - |
| **CC Body** | | | | | | |
| Parameter Estimate | -1.3E-02 | -7.0E-04 | -1.1E-02 | 3.2E-02 | - | - |
| t | -8.348 | -0.096 | -1.494 | 3.664 | - | - |
| 95% CI ($10^{-3}$) | -15.5, -9.6 | -14.9, 13.6 | -24.9, 3.4 | 14.7, 48.9 | - | - |
| p-value | <0.001 | 0.923 | 0.136 | <0.001 | - | - |
| **CC Splenium** | | | | | | |
| Parameter Estimate | -8.7E-03 | -7.7E-03 | -1.9E-02 | 4.6E-02 | - | - |
| t | -7.016 | -1.130 | -3.016 | 6.103 | - | - |
| 95% CI ($10^{-3}$) | -11.2, -6.3 | -21.0, 5.7 | -31.7, -6.7 | 31.3, 61.2 | - | - |
| p-value | <0.001 | 0.259 | 0.003 | <0.001 | - | - |
| **Fornix** | | | | | | |
| Parameter Estimate<br>*EBF*M*Age*<br>*EBF*F*Age*<br>*nEBF*M*Age* | 5.2E-03 | 3.7E-01 | 1.0E-01 | -3.0E-01 | - | -1.1E-02<br>-1.6E-02<br>-6.6E-02 |

| | | | | | | |
|---|---|---|---|---|---|---|
| t<br>EBF*M*Age<br>EBF*F*Age<br>nEBF*M*Age | 0.248 | 3.181 | 0.904 | -2.234 | - | -0.465<br>-0.696<br>-2.819 |
| 95% CI ($10^{-3}$)<br>EBF*M*Age<br>EBF*F*Age<br>nEBF*M*Age | -36.4, 46.8 | 139.8, 596.7 | -120.9, 325.8 | -572.9, -35.8 | - | -57.7, 35.7<br>-62.4, 29.9<br>-113.1, -19.9 |
| p-value<br>EBF*M*Age<br>EBF*F*Age<br>nEBF*M*Age | 0.805 | 0.002 | 0.367 | 0.027 | - | 0.643<br>0.487<br>0.005 |
| **Cingulum** | | | | | | |
| Parameter Estimate | -1.1E-02 | -6.2E-03 | -1.0E-02 | 2.8E-02 | - | - |
| t | -2.090 | -1.156 | -10.989 | 4.755 | - | - |
| 95% CI ($10^{-3}$) | -20.5, -0.6 | -16.9, 4.4 | -12.4, -8.6 | 16.7, 40.2 | - | - |
| p-value | 0.037 | 0.249 | <0.001 | <0.001 | - | - |
| **IFOF** | | | | | | |
| Parameter Estimate | -2.1E-02 | -7.6E-03 | -1.5E-02 | 3.5E-02 | - | - |
| t | -4.588 | -1.482 | -17.094 | 6.332 | - | - |
| 95% CI | -30.2, -12.0 | -17.7, 2.5 | -16.8, -12.6 | 23.8, 45.5 | - | - |
| p-value | <0.001 | 0.139 | <0.001 | <0.001 | - | - |
| **ILF** | | | | | | |
| Parameter Estimate | -1.8E-02 | 1.0E-02 | -1.7E-02 | 3.6E-02 | - | - |
| t | -3.284 | 1.732 | -16.007 | 5.462 | - | - |
| 95% CI | -28.8, -7.2 | -1.4, 21.3 | -19.1, -14.9 | 23.1, 49.1 | - | - |
| p-value | 0.001 | 0.084 | <0.001 | <0.001 | - | - |
| **SLF** | | | | | | |
| Parameter Estimate | -2.0E-02 | -5.0E-04 | -1.1E-02 | 3.2E-02 | - | - |
| t | -4.097 | -0.099 | -12.265 | 5.625 | - | - |
| 95% CI | -29.1, -10.0 | -10.6, 9.6 | -12.6, -8.9 | 20.5, 42.9 | - | - |
| p-value | <0.001 | 0.922 | <0.001 | <0.001 | - | - |
| **Pyramidal Tract** | | | | | | |
| Parameter Estimate<br>EBF*M*Age<br>EBF*F*Age<br>nEBF*M*Age | -6.4E-03 | 1.6E-02 | 1.8E-02 | - | - | -4.2E-0.3<br>-4.1E-0.3<br>-2.9E-0.3 |
| t<br>EBF*M*Age<br>EBF*F*Age<br>nEBF*M*Age | -3.353 | 1.970 | 2.210 | - | - | -1.516<br>-2.267<br>-1.672 |
| 95% CI ($10^{-3}$)<br>EBF*M*Age<br>EBF*F*Age<br>nEBF*M*Age | -10.1, -2.6 | -0.05, 31.2 | 2.0, 34.9 | - | - | -9.7, 1.3<br>-7.7, -0.5<br>-6.4, 0.5 |
| p-value<br>EBF*M*Age<br>EBF*F*Age<br>nEBF*M*Age | <0.001 | 0.051 | 0.028 | - | - | 0.131<br>0.025<br>0.097 |
| **UF** | | | | | | |
| Parameter Estimate | -1.7E-02 | -1.0E-02 | -1.3E-02 | 1.7E-02 | - | - |
| t | -3.776 | -2.109 | -13.802 | 3.187 | - | - |
| 95% CI | -26.4, -8.3 | -20.0, -0.7 | -15.3, -11.5 | 6.4, 27.7 | - | - |
| p-value | <0.001 | 0.036 | <0.001 | 0.002 | - | - |

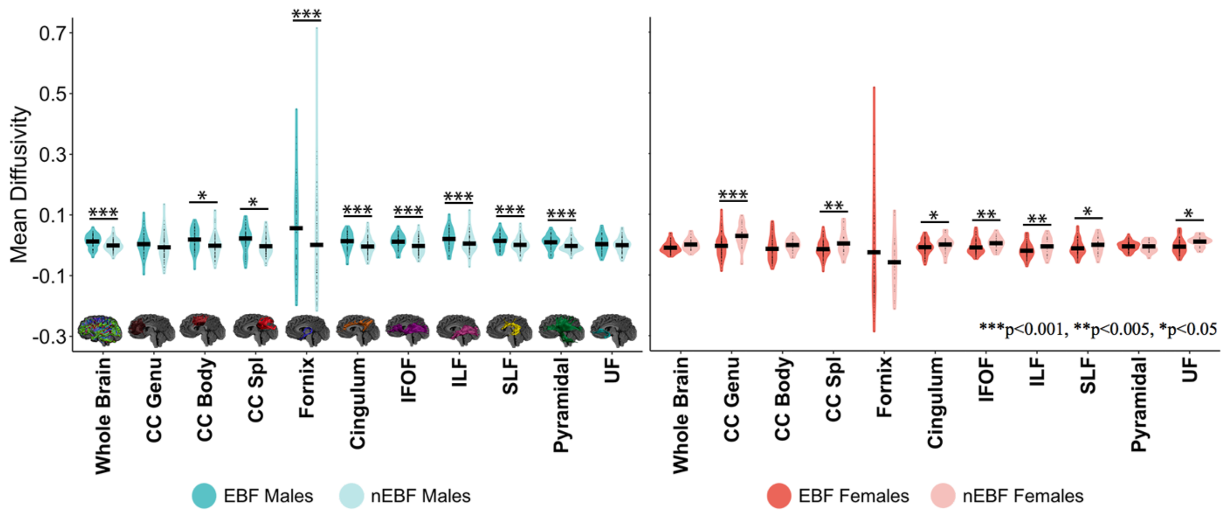


**Supplementary Figure S1.** Breastfeeding group and sex interactions for mean diffusivity in the whole brain and individual white matter tracts. EBF males showed significantly higher MD than nEBF males in the whole brain and most individual tracts, except the genu of the corpus callosum and UF. EBF females had significantly lower MD than nEBF females in most individual tracts, but not the whole brain, the body of the corpus callosum, the fornix and the pyramidal tract. MD values are represented as mean±95% confidence interval and are corrected for age at scan. Data is represented as mean ± 95% confidence interval for FA values for whole brain and individual tracts in EBF males vs. nEBF males and EBF females vs. nEBF females. FA values are corrected for age at the participant's scan. CC Genu = genu of the corpus callosum, CC Body = body of the corpus callosum, CC Splenium = splenium of the corpus callosum, IFOF = inferior fronto-occipital fasciculus, ILF = inferior longitudinal fasciculus, SLF = superior longitudinal fasciculus, UF = uncinate fasciculus.

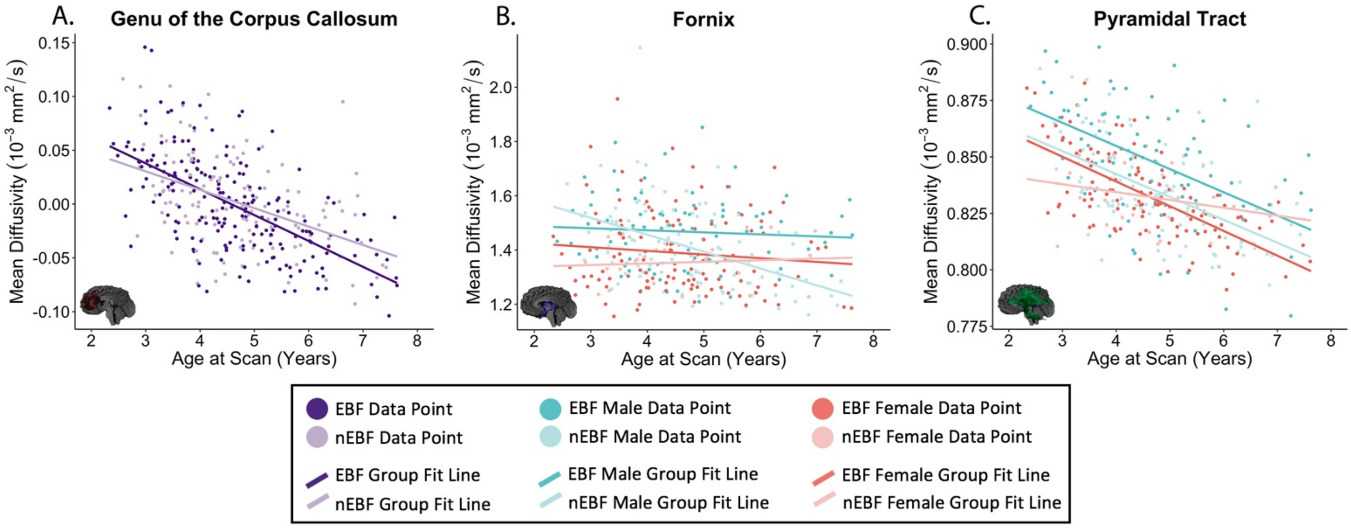


**Supplementary Figure S2.** Significant breastfeeding group-age interaction for mean diffusivity (MD) in the genu of the corpus callosum (F=7.6, p=0.007; A), where the exclusively breastfed group shows steeper decreases in MD. Significant breastfeeding group, age, and sex interactions for mean diffusivity in the fornix (F=6.3, p<0.001; B) and the pyramidal tract (F=3.9, p=0.011; C). MD values for (A) are corrected for sex.

AD: The group-sex interaction is significant in all tracts except for the cingulum, with EBF females < nEBF females and EBF males > nEBF males (Supplementary Table S2). Exceptions occurred in the fornix and the pyramidal tract where nEBF females showed lower AD. The group-age interaction was significant in the genu and body of the corpus callosum, where the EBF group showed steeper age-related decreases for AD in the genu and the nEBF group showed steeper age-related decreases for AD in the body. The group-sex-age interaction was significant in the fornix and the pyramidal tract. All findings survived correction for multiple comparisons. Most findings remained significant after controlling for perinatal and sociodemographic factors, child's FSIQ and motion (number of volumes removed) except for the interactions in the pyramidal tract and the group-age interaction in the body of the corpus callosum.

**Supplementary Table S2. Linear mixed effects results analyzing differences in AD between breastfeeding groups (i.e. EBF vs. nEBF).**

| Axial Diffusivity (AD) | | | | | | |
|---|---|---|---|---|---|---|
| | Main Effects | | | Interactions | | |
| | Group | Sex | Age | Group*Sex | Group*Age | Group*Age*Sex |
| **CC Genu** | | | | | | |
| Parameter Estimate | -1.7E-02 | -2.9E-02 | -1.9E-03 | 4.6E-02 | -8.9E-03 | - |
| t | -5.299 | -3.094 | -0.096 | 4.239 | -2.295 | - |
| 95% CI ($10^{-3}$) | -22.9, -10.4 | -47.6, -10.6 | -42.1, 38.2 | 24.6, 67.3 | -16.7, -1.2 | - |
| p-value | <0.001 | 0.26 | 0.276 | <0.001 | 0.024 | - |
| **CC Body** | | | | | | |
| Parameter Estimate | -1.5E-02 | 1.6E-02 | -4.4E-02 | 2.3E-02 | 7.7E-03 | - |
| t | -5.461 | 1.885 | -2.446 | 2.290 | 2.265 | - |
| 95% CI ($10^{-3}$) | -20.0, -9.4 | -0.7, 32.5 | -80.1, -8.6 | 3.2, 42.4 | 1.0, 14.5 | - |
| p-value | <0.001 | <0.001 | 0.057 | 0.023 | 0.025 | - |
| **CC Splenium** | | | | | | |
| Parameter Estimate | -6.1E-03 | 3.0E-03 | -2.1E-02 | 5.0E-02 | - | - |
| t | -3.667 | 0.322 | -2.395 | 4.923 | - | - |
| 95% CI ($10^{-3}$) | -9.4, -2.8 | -15.1, 21.0 | -37.4, -3.7 | 30.1, 70.3 | - | - |
| p-value | <0.001 | <0.001 | 0.37 | <0.001 | - | - |
| **Fornix** | | | | | | |

| | | | | | | |
|---|---|---|---|---|---|---|
| Parameter Estimate<br>EBF*M*Age<br>EBF*F*Age<br>nEBF*M*Age | 2.0E-02 | 5.1E-01 | 1.7E-01 | -4.5E-01 | - | -1.2E-02<br>-2.5E-0.2<br>-8.7E-02 |
| t<br>EBF*M*Age<br>EBF*F*Age<br>nEBF*M*Age | 0.706 | 3.326 | 1.152 | -2.530 | - | -0.374<br>-0.804<br>-2.804 |
| 95% CI ($10^{-3}$)<br>EBF*M*Age<br>EBF*F*Age<br>nEBF*M*Age | -35.3, 74.7 | 207.6, 813.1 | -122.1, 465.1 | -803.6, -99.6 | - | -73.1, 49.9<br>-8.5, 3.6<br>-14.9, -2.6 |
| p-value<br>EBF*M*Age<br>EBF*F*Age<br>nEBF*M*Age | 0.219 | 0.002 | 0.544 | 0.012 | - | 0.709<br>0.422<br>0.006 |
| **Cingulum** | | | | | | |
| Parameter Estimate | -8.3E-03 | 1.2E-02 | 1.9E-02 | - | - | - |
| t | -5.236 | 4.001 | 4.333 | - | - | - |
| 95% CI ($10^{-3}$) | -16.5, -3.0E-02 | -626524.6, 626549.1 | 5.9, 32.2 | - | - | - |
| p-value | 0.05 | 0.78 | 0.017 | - | - | - |
| **IFOF** | | | | | | |
| Parameter Estimate | -1.1E-02 | -3.7E-03 | -6.1E-03 | 2.2E-02 | - | - |
| t | -7.227 | -0.483 | -0.845 | 2.527 | - | - |
| 95% CI ($10^{-3}$) | -13.5, -7.7 | -18.9, 11.5 | -20.3, 8.1 | 4.9, 39.4 | - | - |
| p-value | <0.001 | 0.094 | 0.257 | 0.012 | - | - |
| **ILF** | | | | | | |
| Parameter Estimate | -1.1E-02 | 1.5E-02 | -1.9E-02 | 3.5E-02 | - | - |
| T | -8.812 | 2.231 | -2.792 | 4.457 | - | - |
| 95% CI ($10^{-3}$) | -13.4, -8.5 | 1.8, 29.1 | -32.0, -5.5 | 19.7, 50.8 | - | - |
| p-value | <0.001 | <0.001 | 0.775 | <0.001 | - | - |
| **SLF** | | | | | | |
| Parameter Estimate | -8.6E-03 | -3.9E-04 | -5.8E-03 | 2.58E-02 | - | - |
| t | -7.279 | -0.061 | -0.964 | 3.531 | - | - |
| 95% CI ($10^{-3}$) | -10.9, -6.2 | -13.0, 12.3 | -17.6, 6.0 | 11.4, 40.2 | - | - |
| p-value | <0.001 | 0.001 | 0.052 | <0.001 | - | - |
| **Pyramidal Tract** | | | | | | |
| Parameter Estimate<br>EBF*M*Age<br>EBF*F*Age<br>nEBF*M*Age | -1.7E-03 | 5.8E-02 | 4.5E-02 | -7.2E-02 | - | 5.0E-04<br>-7.5E-03<br>-7.3E-03 |
| t<br>EBF*M*Age<br>EBF*F*Age<br>nEBF*M*Age | -0.437 | 2.751 | 2.383 | -3.106 | - | 0.116<br>-1.934<br>-1.697 |
| 95% CI ($10^{-3}$)<br>EBF*M*Age<br>EBF*F*Age<br>nEBF*M*Age | -9.3, 5.9 | 16.5, 99.7 | 7.7, 81.7 | -118.1, -26.4 | - | -8.0, 9.1<br>-15.1, 0.1<br>-15.9, 1.2 |
| p-value<br>EBF*M*Age<br>EBF*F*Age<br>nEBF*M*Age | <0.001 | 0.06 | 0.46 | 0.002 | - | 0.908<br>0.055<br>0.091 |

| | | | | | | |
|---|---|---|---|---|---|---|
| **UF** | | | | | | |
| Parameter Estimate | -1.0E-02 | 3.4E-02 | -1.0E-02 | 1.7E-02 | - | - |
| t | -6.195 | 2.605 | -1.628 | 2.335 | - | - |
| 95% CI ($10^{-3}$) | -13.7, 7.0 | 8.3, 60.2 | -0.22.1, 2.1 | 2.7, 31.9 | - | - |
| p-value | <0.001 | 0.001 | 0.719 | 0.021 | - | - |

RD: The group-sex interaction was significant for RD in all tracts (Supplementary Table S3), with EBF females < nEBF females and EBF males > nEBF males. An exception occurred in the fornix where nEBF females showed lower RD. The group-age interaction was significant for the IFOF and the pyramidal tract, where the EBF group shows steeper age-related decreases for RD in both tracts. All findings survived correction for multiple comparisons except for the group-sex interaction in the fornix. All findings remained significant after controlling for perinatal and sociodemographic factors, child's FSIQ and motion (number of volumes removed).

**Supplementary Table S3. Linear mixed effects results analyzing differences in RD between breastfeeding groups (i.e. EBF vs. nEBF).**

| | **Radial Diffusivity (RD)** | | | | | |
|---|---|---|---|---|---|---|
| | **Main Effects** | | | **Interactions** | | |
| | **Age** | **Sex** | **Group** | **Group*Sex** | **Group*Age** | **Group*Age*Sex** |
| **CC Genu** | | | | | | |
| Parameter Estimate | -2.0E-02 | -4.1E-02 | -3.4E-02 | 4.8E-02 | - | - |
| t | -13.717 | -5.251 | -4.749 | 5.475 | - | - |
| 95% CI ($10^{-3}$) | -23.2, -17.4 | -56.8, -25.8 | -47.5, -19.7 | 30.5, 64.8 | - | - |
| p-value | <0.001 | <0.001 | 0.027 | <0.001 | - | - |
| **CC Body** | | | | | | |
| Parameter Estimate | -1.4E-02 | -8.6E-03 | -1.3E-02 | 3.6E-02 | - | - |
| t | -8.682 | -1.158 | -1.756 | 3.999 | - | - |
| 95% CI ($10^{-3}$) | -16.8, -10.6 | -23.3, 6.0 | -27.6, 1.6 | 18.2, 53.5 | - | - |
| p-value | <0.001 | 0.039 | 0.273 | <0.001 | - | - |
| **CC Splenium** | | | | | | |
| Parameter Estimate | -9.8E-03 | -1.3E-02 | -1.9E-02 | 4.5E-02 | - | - |
| t | -8.345 | -1.944 | -3.068 | 6.201 | - | - |
| 95% CI ($10^{-3}$) | -12.2, -7.5 | -25.4, 0.2 | -30.6, -6.6 | 30.6, 59.2 | - | - |
| p-value | <0.001 | 0.007 | 0.29 | <0.001 | - | - |
| **Fornix** | | | | | | |
| Parameter Estimate<br>*EBF*M*Age*<br>*EBF*F*Age*<br>*nEBF*M*Age* | -2.2E-03 | 3.0E-01 | 7.0E-02 | -2.4E-01 | - | <br>-9.2E-03<br>-1.3E-02<br>-5.6E-0.2 |

| | | | | | | |
|---|---|---|---|---|---|---|
| t<br>EBF*M*Age<br>EBF*F*Age<br>nEBF*M*Age | -0.120 | 2.984 | 0.718 | -2.059 | - | -0.448<br>-0.632<br>-2.746 |
| 95% CI ($10^{-3}$)<br>EBF*M*Age<br>EBF*F*Age<br>nEBF*M*Age | -38.1, 33.7 | 100.5, 492.8 | -122.3, 262.3 | -473.3, -10.3 | - | -49.4, 31.1<br>-52.5, 27.0<br>-95.9,-15.7 |
| p-value<br>EBF*M*Age<br>EBF*F*Age<br>nEBF*M*Age | <0.001 | 0.003 | 0.387 | 0.041 | - | <0.001<br>0.528 |
| **Cingulum** | | | | | | |
| Parameter Estimate | -1.5E-02 | -1.5E-02 | -2.3E-02 | 4.0E-02 | - | - |
| t | -220.172 | -2.855 | -4.946 | 7.604 | - | - |
| 95% CI ($10^{-3}$) | -14.9, -14.7 | -25.4, -4.7 | -32.7, -14.1 | 29.7, 50.4 | - | - |
| p-value | <0.001 | 0.059 | 0.209 | <0.001 | - | - |
| **IFOF** | | | | | | |
| Parameter Estimate | -1.1E-02 | 9.4E-03 | 4.8E-03 | 3.7E-02 | -5.5E-03 | - |
| t | -5.305 | 1.003 | 0.486 | 7.376 | -2.854 | - |
| 95% CI ($10^{-3}$) | -15.0, -6.9 | -9.1, 28.0 | -14.8, 24.5 | 26.9, 46.4 | -9.3, -1.7 | - |
| p-value | <0.001 | <0.001 | 0.013 | <0.001 | 0.005 | - |
| **ILF** | | | | | | |
| Parameter Estimate | -2.0E-02 | 6.9E-03 | -2.1E-02 | 3.9E-02 | - | - |
| t | -17.472 | 1.125 | -3.829 | 5.849 | - | - |
| 95% CI ($10^{-3}$) | -22.0, -17.6 | -5.1, 18.9 | -31.7, -10.2 | 26.2, 52.7 | - | - |
| p-value | <0.001 | <0.001 | 0.726 | <0.001 | - | - |
| **SLF** | | | | | | |
| Parameter Estimate | -1.2E-02 | -6.7E-04 | -2.2E-02 | 3.4E-02 | - | - |
| t | -13.563 | -0.128 | -4.626 | 6.539 | - | - |
| 95% CI ($10^{-3}$) | -13.6, -10.1 | -10.9, 9.6 | -31.3, -12.6 | 23.8, 44.4 | - | - |
| p-value | <0.001 | <0.001 | 0.068 | <0.001 | - | - |
| **Pyramidal Tract** | | | | | | |
| Parameter Estimate | -9.2E-03 | -6.9E-03 | 1.8E-02 | 2.0E-02 | -5.1E-03 | - |
| t | -6.568 | -1.699 | 2.061 | 4.328 | -3.085 | - |
| 95% CI ($10^{-3}$) | -11.9, -6.4 | -14.8, 1.1 | 0.7, 35.2 | 10.7, 28.7 | -8.4, -1.8 | - |
| p-value | <0.001 | 0.189 | 0.001 | <0.001 | 0.003 | - |
| **UF** | | | | | | |
| Parameter Estimate | -1.4E-02 | 2.1E-02 | -2.3E-02 | 3.0E-02 | - | - |
| t | -8.666 | 1.784 | -4.244 | 4.610 | - | - |
| 95% CI ($10^{-3}$) | -17.8, -11.1 | -2.3, 44.9 | -34.1, -12.5 | 17.4, 43.4 | - | - |
| p-value | <0.001 | 0.001 | 0.014 | <0.001 | - | - |

*S3.2 Breastfeeding duration analysis*

MD: The duration-sex interaction was significant for MD in the body of the corpus callosum ($p<0.001$) and the IFOF ($p=0.019$) (Supplementary Table S4). In both tracts, increases in MD were associated with a longer breastfeeding duration for males; additionally, only in the body of the corpus callosum, decreases in MD were associated with a longer breastfeeding duration for females (Supplementary Figure S3A). There was a main effect of duration for MD in the whole brain as well as the following individual tracts: the fornix, the cingulum, the ILF, the SLF, and the pyramidal tract where increases in MD were associated with a longer breastfeeding duration both globally and within each tract (Supplementary Figure S3B, S3C). Findings in the ILF and the IFOF did not survive correction for multiple comparisons. All findings remained significant after controlling for perinatal/sociodemographic covariates, child's FSIQ and motion (number of volumes removed) except for the main effect in the whole brain and the cingulum which did not survive correction for child's FSIQ.

**Supplementary Table S4. Linear mixed effects results to analyze associations between MD and total duration of breastfeeding.**

| | **Mean Diffusivity (MD)** | | | |
|---|---|---|---|---|
| | **Main Effects** | | | **Interaction** |
| | **Age** | **Sex** | **Duration** | **Duration*Sex** |
| **Whole Brain** | | | | |
| Parameter Estimate | -1.3E-02 | 1.5E-02 | 3.8E-03 | - |
| t | -16.920 | 20.509 | 2.718 | - |
| 95% CI ($10^{-3}$) | -14.7, -11.6 | 13.7, 16.6 | 1.0, 6.5 | - |
| p-value | <0.001 | <0.001 | 0.007 | - |
| **CC Genu** | | | | |
| Parameter Estimate | -2.1E-02 | -7.1E-03 | -1.5E-03 | - |
| t | -13.461 | -2.004 | -0.452 | - |
| 95% CI ($10^{-3}$) | -24.5, -18.2 | -14.7, 0.6 | -7.8, 4.9 | - |
| p-value | <0.001 | 0.067 | 0.651 | - |
| **CC Body** | | | | |
| Parameter Estimate | -1.2E-02 | -9.2E-03 | -1.1E-02 | 2.1E-02 |
| t | -7.584 | -1.073 | -2.187 | 3.390 |
| 95% CI ($10^{-3}$) | -14.9, -8.7 | -26.2, 7.7 | -21.1, -1.1 | 8.9, 33.5 |
| p-value | <0.001 | 0.284 | 0.030 | <0.001 |
| **CC Splenium** | | | | |
| Parameter Estimate | -9.1E-03 | 2.1E-02 | 4.8E-03 | - |
| t | -6.419 | 6.951 | 1.738 | - |
| 95% CI ($10^{-3}$) | -12.0, -6.3 | 14.7, 27.1 | -0.7, 10.4 | - |
| p-value | <0.001 | <0.001 | 0.085 | - |
| **Fornix** | | | . | |
| Parameter Estimate | -2.2E-02 | 6.1E-02 | 5.3E-02 | - |
| t | -3.603 | 4.143 | 4.561 | - |
| 95% CI ($10^{-3}$) | -34.2, -10.0 | 32.2, 90.4 | 30.2, 76.1 | - |
| p-value | <0.001 | <0.001 | <0.001 | - |
| **Cingulum** | | | | |
| Parameter Estimate | -1.1E-02 | 1.2E-02 | 5.8E-03 | - |

| | | | | |
|---|---|---|---|---|
| t | -10.724 | 5.495 | 2.710 | - |
| 95% CI ($10^{-3}$) | -13.0, -9.0 | 7.6, 16.8 | 1.6, 10.0 | - |
| p-value | <0.001 | <0.001 | 0.007 | - |
| **IFOF** | | | | |
| Parameter Estimate | -1.5E-02 | 3.1E-05 | -4.8E-03 | 1.0E-02 |
| t | -14.170 | 0.005 | -1.351 | 2.354 |
| 95% CI ($10^{-3}$) | -16.7, -12.6 | -11.8, 11.9 | -11.7, 2.2 | 1.7, 18.6 |
| p-value | <0.001 | 0.996 | 0.178 | 0.019 |
| **ILF** | | | | |
| Parameter Estimate | -1.7E-02 | 3.3E-02 | 4.8E-03 | - |
| t | -15.465 | 17.076 | 2.072 | - |
| 95% CI ($10^{-3}$) | -19.6, -15.1 | 29.6, 37.3 | 0.2, 9.4 | - |
| p-value | <0.001 | <0.001 | 0.039 | - |
| **SLF** | | | | |
| Parameter Estimate | -1.1E-02 | 1.9E-02 | 4.0E-03 | - |
| t | -11.086 | 8.492 | 2.063 | - |
| 95% CI ($10^{-3}$) | -13.0, -9.0 | 14.5, 23.6 | 0.2, 7.8 | - |
| p-value | 0.04 | <0.001 | <0.001 | - |
| **Pyramidal Tract** | | | | |
| Parameter Estimate | -9.8E-03 | 8.4E-03 | 5.5E-03 | - |
| t | -12.691 | 5.078 | 3.551 | - |
| 95% CI ($10^{-3}$) | -11.3, -8.3 | 5.1, 11.7 | 2.4, 8.5 | - |
| p-value | <0.001 | <0.001 | <0.001 | - |
| **UF** | | | | |
| Parameter Estimate | -1.3E-02 | 3.9E-03 | 2.6E-04 | - |
| t | -13.112 | 1.928 | 0.138 | - |
| 95% CI ($10^{-3}$) | -14.9, -10.9 | -1.0, 8.8 | -3.4, 4.0 | - |
| p-value | <0.001 | 0.101 | 0.891 | - |

If interactions were not significant, they were removed from the model.

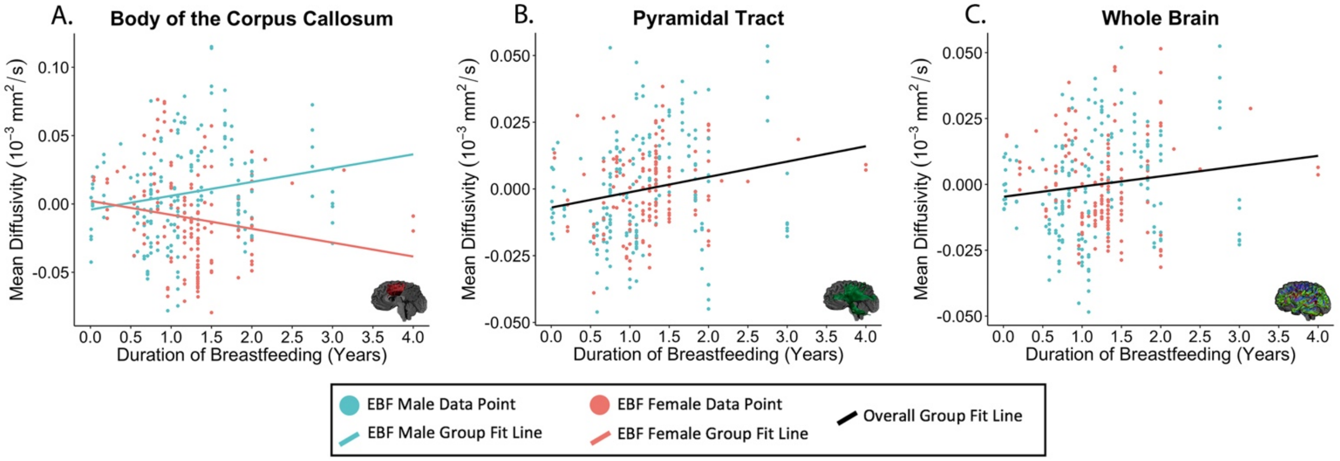


**Supplementary Figure S3.** Significant interactions were noted between breastfeeding duration and child's sex for MD in the body of the corpus callosum (F=11.5, p<0.001; A). A significant main effect of breastfeeding duration was noted for MD in the pyramidal tract (F=161.1, p<0.001; B) and the whole brain (F=295.1, p=0.007; C), where longer breastfeeding was associated with higher MD. Plot B and C are representative of the main effect also observed in the fornix, the cingulum, and the SLF. MD values are corrected for age at participant's scan.

AD: The duration-sex interaction was significant for AD in the body of the corpus callosum, ILF and the UF (Supplementary Table S5). In all tracts, higher AD was associated with a longer breastfeeding duration for males; additionally, only in the body of the corpus callosum, lower AD was associated with a longer breastfeeding duration for females. There was also a main effect of duration for AD in the fornix and the cingulum, where higher AD was associated with longer durations of breastfeeding in both tracts. All findings survived correction for multiple comparisons except for the interaction effect in the UF. All findings remained significant after controlling for perinatal and sociodemographic covariates, child's FSIQ and motion (number of volumes removed).

**Supplementary Table S5. Linear mixed effects results to analyze associations between AD and total duration of breastfeeding.**

| | Axial Diffusivity | | | |
|---|---|---|---|---|
| | Main Effects | | | Interactions |
| | Age | Sex | Duration | Duration*Sex |
| **CC Genu** | | | | |
| Parameter Estimate | -2.3E-02 | 9.3E-03 | -5.4E-03 | - |
| t | -12.736 | 2.682 | -1.409 | - |
| 95% CI ($10^{-3}$) | -26.3, -19.2 | 2.4, 16.2 | -13.0, 2.2 | - |
| p-value | <0.001 | 0.008 | 0.160 | - |
| **CC Body** | | | | |
| Parameter Estimate | -9.5E-03 | 1.7E-03 | -1.2E-02 | 2.2E-02 |
| t | -5.605 | 0.175 | -1.964 | 3.056 |
| 95% CI ($10^{-3}$) | -12.9, -6.2 | -17.7, 21.2 | -23.1, 3722.8 | 7.8, 36.0 |
| p-value | <0.001 | 0.861 | 0.871 | 0.002 |
| **CC Splenium** | | | | |
| Parameter Estimate | -6.7E-03 | 3.4E-02 | 5.3E-03 | - |
| t | -3.626 | 8.832 | 1.459 | - |
| 95% CI ($10^{-3}$) | -10.3, -3.0 | 26.2, 41.3 | -1.8, 12.4 | - |
| p-value | <0.001 | <0.001 | 0.146 | - |
| **Fornix** | | | | |
| Parameter Estimate | -1.7E-02 | 9.2E-02 | 7.4E-02 | - |
| t | -2.148 | 5.007 | 4.885 | - |
| 95% CI ($10^{-3}$) | -32.7, -1.4 | 55.6, 128.5 | 44.4, 104.4 | - |
| p-value | 0.033 | <0.001 | <0.001 | - |
| **Cingulum** | | | | |
| Parameter Estimate | -7.8E-03 | 1.0E-02 | 1.3E-02 | - |
| t | -4.336 | 2.409 | 3.557 | - |
| 95% CI ($10^{-3}$) | -11.3, -4.2 | 1.7, 18.7 | 5.6, 19.6 | - |

| | | | | |
|---|---|---|---|---|
| p-value | <0.001 | 0.02 | <0.001 | - |
| **IFOF** | | | | |
| Parameter Estimate | -1.0E-02 | 8.1E-03 | 4.6E-03 | - |
| t | -6.993 | 2.454 | 1.486 | - |
| 95% CI ($10^{-3}$) | -13.5, -7.5 | 1.5, 14.7 | -1.5, 10.6 | - |
| p-value | <0.001 | 0.016 | 0.139 | - |
| **ILF** | | | | |
| Parameter Estimate | -1.0E-02 | 1.1E-02 | -8.1E-03 | 2.0E-02 |
| t | -7.661 | 1.370 | -1.670 | 3.505 |
| 95% CI ($10^{-3}$) | -12.7, -7.5 | -4.8, 26.7 | -17.6, 1.4 | 8.9, 31.8 |
| p-value | <0.001 | 0.172 | 0.468 | <0.001 |
| **SLF** | | | | |
| Parameter Estimate | -8.3E-03 | 1.3E-02 | 3.4E-03 | - |
| t | -7.041 | 5.051 | 1.310 | - |
| 95% CI ($10^{-3}$) | -10.7, -5.9 | 8.0, 18.7 | -1.7, 8.4 | - |
| p-value | <0.001 | <0.001 | 0.192 | - |
| **Pyramidal Tract** | | | | |
| Parameter Estimate | -5.5E-03 | 1.8E-02 | 3.8E-03 | - |
| t | -5.026 | 7.781 | 1.787 | - |
| 95% CI ($10^{-3}$) | -7.6, -3.3 | 13.6, 23.0 | -0.4, 7.9 | - |
| p-value | <0.001 | <0.001 | 0.075 | - |
| **UF** | | | | |
| Parameter Estimate | -1.3E-02 | 6.5E-03 | -3.7E-03 | 1.2E-02 |
| t | -10.667 | 0.867 | -0.814 | 2.217 |
| 95% CI ($10^{-3}$) | -15.4, -10.5 | -8.3, 21.4 | -12.6, 5.2 | 1.4, 23.2 |
| p-value | <0.001 | 0.387 | 0.377 | 0.027 |

RD: The duration-sex interaction was significant for RD in the body of the corpus callosum, cingulum and the IFOF (Supplementary Table S6). In the body of the corpus callosum and the cingulum, increases in RD for males and decreases in RD for females were associated with longer durations of breastfeeding. There was also a main effect of duration for RD in the following tracts: the splenium of the corpus callosum, the fornix, the SLF, and the pyramidal tract. For all tracts, increases in RD were associated with longer durations of breastfeeding. All findings survived correction for multiple comparisons except for the main effect in the splenium and the interaction effect in the IFOF. All findings remained significant after controlling for perinatal and sociodemographic factors, child’s FSIQ and motion (number of volumes removed) except for the main effect in the SLF which did not remain significant after accounting for child’s FSIQ.

**Supplementary Table S6. Linear mixed effects results to analyze associations between RD and total duration of breastfeeding.**

| | Radial Diffusivity | | | |
|---|---|---|---|---|
| | Main Effects | | | Interactions |
| | Age | Sex | Duration | Duration*Sex |
| **CC Genu** | | | | |
| Parameter Estimate | -2.0E-02 | -1.0E-02 | -9.7E-04 | - |
| t | -12.685 | -2.759 | -0.306 | - |
| 95% CI ($10^{-3}$) | -23.7, -17.2 | -17.6, -2.4 | -7.3, 5.3 | - |
| p-value | <0.001 | 0.012 | 0.760 | - |
| **CC Body** | | | | |
| Parameter Estimate | -1.3E-02 | -1.4E-02 | -1.1E-02 | 2.1E-02 |
| t | -7.902 | -1.601 | -2.105 | 3.219 |
| 95% CI ($10^{-3}$) | -16.3, -9.8 | -31.8, 3.3 | -21.3, -0.7 | 8.1, 33.5 |
| p-value | <0.001 | 0.111 | 0.852 | 0.001 |
| **CC Splenium** | | | | |
| Parameter Estimate | -1.1E-02 | 1.6E-02 | 5.4E-03 | - |
| t | -8.182 | 7.527 | 2.052 | - |
| 95% CI ($10^{-3}$) | -13.1, -8.0 | 12.0, 20.6 | 0.2, 10.5 | - |
| p-value | <0.001 | <0.001 | 0.041 | - |
| **Fornix** | | | | |
| Parameter Estimate | -2.5E-02 | 4.7E-02 | 4.2E-02 | - |
| t | -4.642 | 3.620 | 4.135 | - |
| 95% CI ($10^{-3}$) | -35.2, -14.2 | 21.4, 72.3 | 21.8, 61.4 | - |
| p-value | <0.001 | <0.001 | <0.001 | - |
| **Cingulum** | | | | |
| Parameter Estimate | -1.4E-02 | -4.0E-03 | -7.6E-03 | 1.5E-02 |
| t | -13.814 | -0.625 | -1.977 | 3.369 |
| 95% CI ($10^{-3}$) | -16.9, -11.8 | -16.6, 8.6 | -15.1, -2.3E-02 | 6.4, 24.3 |
| p-value | <0.001 | 0.532 | 0.960 | <0.001 |
| **IFOF** | | | | |
| Parameter Estimate | -1.7E-02 | 1.5E-03 | -4.8E-03 | 1.0E-02 |
| t | -16.867 | 0.258 | -1.442 | 2.432 |
| 95% CI ($10^{-3}$) | -18.7, -14.8 | -9.8, 12.7 | -11.4, 1.8 | 1.9, 18.0 |
| p-value | <0.001 | 0.797 | 0.942 | 0.016 |
| **ILF** | | | | |
| Parameter Estimate | -2.0E-02 | 2.9E-02 | 4.1E-03 | - |
| t | -16.525 | 10.270 | 1.641 | - |
| 95% CI ($10^{-3}$) | -22.8, -17.9 | 23.2, 35.5 | -0.8, 9.0 | - |
| p-value | <0.001 | <0.001 | 0.102 | - |
| **SLF** | | | | |
| Parameter Estimate | -1.2E-02 | 2.2E-02 | 5.2E-03 | - |
| t | -11.847 | 9.139 | 2.577 | - |
| 95% CI ($10^{-3}$) | -14.1, -10.1 | 17.3, 27.1 | 1.2, 9.2 | - |
| p-value | <0.001 | <0.001 | 0.011 | - |
| **Pyramidal Tract** | | | | |
| Parameter Estimate | -1.3E-02 | 6.5E-03 | 6.9E-03 | - |
| t | -20.345 | 4.391 | 4.305 | - |
| 95% CI ($10^{-3}$) | -14.5, 11.9 | 3.5, 9.4 | 3.7, 10.0 | - |
| p-value | <0.001 | <0.001 | <0.001 | - |
| **UF** | | | | |
| Parameter Estimate | -1.5E-02 | 4.1E-02 | 1.3E-03 | - |
| t | -8.241 | 3.666 | 0.580 | - |

| | | | | |
|---|---|---|---|---|
| 95% CI ($10^{-3}$) | -18.0, -11.0 | 19.1, 63.6 | -3.2, 5.8 | - |
| p-value | <0.001 | <0.001 | 0.563 | - |
